# Phytoplankton and giant virus dynamics during different monsoon seasons, a fish kill, and a toxic bloom in a eutrophic mariculture area

**DOI:** 10.1101/2025.05.12.653504

**Authors:** Andrian P. Gajigan, Gianina Cassandra May B. Apego, Eunice Lois D. Gianan, Rachel B. Francisco, Jun Jet Tai, Garry A. Benico, Charina Lyn A. Repollo, Cesar L. Villanoy, Aletta T. Yñiguez, Cecilia Conaco, Grieg F. Steward

## Abstract

Harmful algal blooms (HABs) pose a significant public health concern and can cause severe economic losses. In Bolinao, Philippines, intensive mariculture has led to diatom- or dinoflagellate-dominated HABs since the 2000s, raising risks of paralytic shellfish poisonings and fish kills. In the context of ongoing HAB monitoring and mitigation efforts, we investigated the phytoplankton and associated giant viruses during two seasons (Nov-Dec 2021 and Apr-May 2022). Our sampling encompassed a saxitoxin episode lasting 2.5 months (April-June 2022), and a fish kill (May 15-16, 2022). We used a low-cost flow-through camera (PlanktoScope) and 18S rRNA gene amplicon sequencing to identify the dominant phytoplankton. We detected the presence of *Alexandrium* sp., a known saxitoxin-producing alga, coincident with a toxin alert that resulted in a shellfish ban, even though it was not the dominant taxon. Using electron microscopy, we observed diverse morphologies of virus-like particles (VLPs), including “giant” icosahedral VLPs (~200 nm capsids). We assembled giant virus genomes from metagenomic data collected at different bloom phases. Phylogenetic and functional analyses suggest that the recovered genomes are primarily from viruses in the orders *Imitervirales* and *Algavirales*, encoding diverse auxiliary metabolic genes involved in nutrient transport and carbohydrate metabolism. The diversity and spatiotemporal dynamics of plankton and their viruses described in this study contributed to our understanding of the broader microbial context of HABs.

**Importance:** HABs are widespread phenomena that can be detrimental to livelihoods and health in coastal communities. Understanding what drives their formation, maintenance, and crash could aid in predicting, and potentially alleviating these events. Here, we report on the diversity and temporal dynamics of phytoplankton and giant viruses during a HAB event. Our findings revealed a diverse array of giant viruses belonging to thirteen families in a eutrophic coastal environment. We found evidence of associations between several phytoplankton-giant virus pairs and found a temporal/seasonal influence on phytoplankton-giant virus community structure. While future research is needed to establish definitive links between the giant viruses and HABs species succession, this study lays the groundwork for future mechanistic studies.

## Introduction

Harmful algal blooms (HABs), exacerbated by intensive fish farming, are a recurring problem in the coastal waters of Bolinao, NW Philippines. The Bolinao mariculture of milkfish (*Chanos chanos*) started in 1970 and intensified in 1995 ^1^. The proliferation of fish pens restricts circulation which, together with nutrient and organic loading from the use of excessive fish feed, contributes to eutrophication that fuels HABs since 2000s ^2–5^. Microbial respiration of organic matter from excess fish feed and crashing phytoplankton blooms contributes to severe oxygen depletion. Stratification resulting from freshwater runoff and surface warming can aggravate the problem by restricting the mixing of oxygen into waters below the thin surface layer, resulting in hypoxia and fish kills. Fish kills in Bolinao have been frequent, with the most notable events occurring in 2002 and 2010 ^6–8^. The 2002 *Prorocentrum minimum* bloom lasted five days, resulting $9 million loss due to fish kill ^6^. Dissolved oxygen (DO) was < 2mg/L, far from the standard ~5-6 mg/L^3^. In response, the local government enforced restrictions on the number of fish structures; however, nutrient concentrations remained high ^9^. In 2010, blooms of *Alexandrium* sp. and *Skeletonema costatum* lasted two months reducing DO to < 0.5 mg L^−1^. This cycle of eutrophication, HABs, hypoxia, and fish kills continues to be a problem, and, in 2018, Bolinao suffered a $2 million loss from another fish kill ^10^. The recurrent HABs provides an opportunity to examine phytoplankton and viral dynamics during these devastating bloom events.

Viruses, which infect diverse phytoplankton^11^, grazers ^12^, and bacteria ^13^ can influence HAB dynamics both directly and indirectly. Viruses that infect HAB-forming species may act directly to suppress initiation or cause termination of a HAB. Viruses are major contributors to phytoplankton mortality ^14^, often causing loss rates similar in magnitude to zooplankton grazing^15,16^. In some cases, viruses may be the primary cause of abrupt bloom termination ^17^. Lysis of a dominant phytoplankton population drives algal species successions ^18,19^ and the concomitant release of dissolved organic material alters the composition and activity of the heterotrophic microbial food web ^20,21^. Viral infections could also indirectly favor HAB blooms by lysing a dominant non-HAB phytoplankton and reducing resource competition. For HAB species adept at using organic nutrients, blooms might be promoted by organic nutrients released by viral lysis ^22^. Thus, viruses may stimulate bloom formation by top-down (control on predators) or bottom-up (increasing nutrient supply) mechanisms. Despite the potentially significant and multifaceted contributions of viruses to bloom ecology, no studies have examined the viral communities associated with Bolinao HABs.

This study focuses on large dsDNA viruses (Phylum *Nucleocytoviricota*), a group of viruses also referred to as Nucleocytoplasmic Large DNA Viruses (NCLDV). There are many smaller RNA- or DNA-containing viruses that could also have a significant influence on HAB dynamics, but we found the NCLDVs, and members of the *Imitervirales* in particular, to be of interest because they are abundant, exceptionally diverse ^23^, and have large genomes with extensive metabolic genes^24,25^. Using environmental sequencing, plankton imaging via low-cost device (PlanktoScope ^26^) and conventional microscopy, we documented phytoplankton and associated giant viruses dynamics across seasons and a saxitoxin HAB episode. This research provides context for future studies and contributes to ongoing efforts to create an ecological model and a HAB early-warning system for the area.

## Materials and methods

### Study sites and sampling

The study was conducted in Guiguiwanen Channel, Bolinao, with samples collected at six stations (**Figure 1**). Two stations were selected to leverage historical and on-going efforts: Station 1_CCMS (16.3868°N 119.9252°E) is the site of a floating platform (Continuous and Comprehensive Monitoring System) established in 2011 within an area densely packed with fish pens ^9^, and Station 2_BML (16.3815°N 119.9125°E) is the Bolinao Marine Laboratory monitoring station established in 1995.

**Figure 1.**
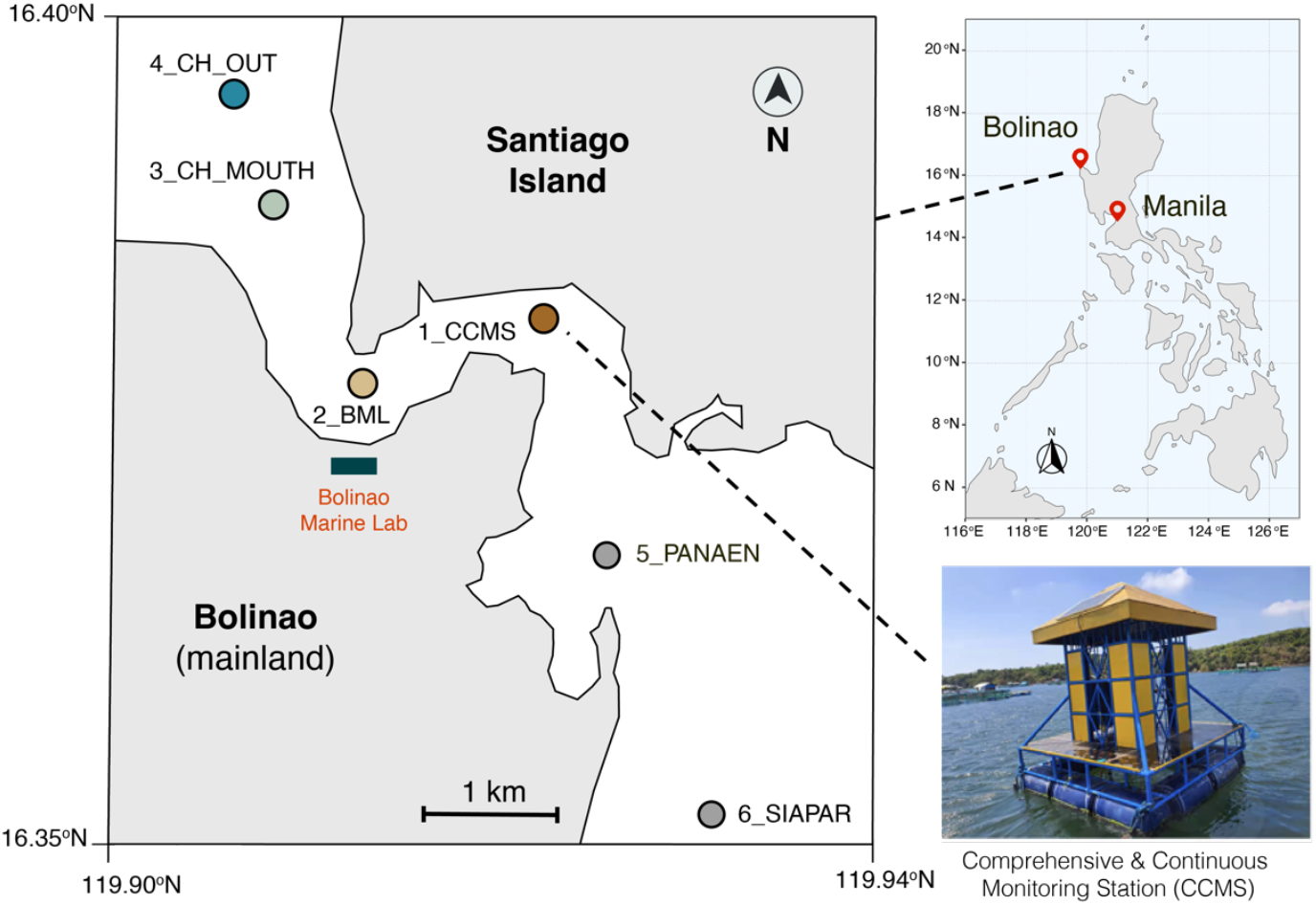
Bolinao map and sampling sites. Bolinao is situated in NW Philippines. We focused our sampling on the Guiguiwanen channel, covering mariculture sites (Stations 1 and 2) and relatively offshore sites (Stations 3 and 4). We sampled inward the channel (5 and 6) whenever logistics and time permits.

Additional stations include Station 3_CH_MOUTH (16.39538°N 119.9052°E) at the northern mouth of the channel and Station 4_CH_OUT (16.4038°N 119.9022°E) even further north. These latter stations correspond, respectively, to the stations Guig2 and Guig1in a previous study ^27^. Hydrographic data were routinely collected at these four stations, but two additional stations further south, within less dense fish pens (**Figure S1)** were also sampled on occasion: Station 5_Panaen and Station 6_Siapar. A Hanna multiparameter probe (HI98194) was deployed at 1m depth to measure temperature, salinity, DO, total dissolved solids (TDS), and pH. Sampling occurred in November-December 2021 and April-May 2022, when HABs are most likely to occur ^28^. The first sampling was done during the Northeast (NE) monsoon (typically October to March), while the second sampling corresponds to the transition period (April) and the onset of the Southwest (SW) monsoon (typically starts mid-May) ^29,30^. A Gratuitous Permit (#0228-22) and Export Clearance (#2022-54 and 55) were secured from the Philippines’ Bureau of Fisheries and Aquatic Resources (BFAR).

### Phytoplankton and virus collection

Phytoplankton samples were collected using a 30-cm wide, 10-μm mesh plankton net, vertically towed from 5 meters to the surface. Discrete seawater samples (~1 m depth) for sequencing and chlorophyll-a analyses were obtained using a 5 L Niskin bottle. Biomass for 18S rRNA gene sequencing was collected by filtration of ~800 mL of seawater onto 0.2 µm polycarbonate filters (Cytiva). Material for metagenomic analysis of the total community, including viruses, was collected by filtration of ~370 mL of seawater on 0.02 µm aluminum oxide filters (Anotop, Cytiva). Pre-filtration through 200 µm nylon mesh was performed for all samples before PlanktoScope imaging, microscopy and biomass collection. Phytoplankton were counted within 2 hours after collection using the PlanktoScope (live imaging) and microscopy (preserved in Lugol’s iodine).

### Chlorophyll-a measurement

Seawater from each site was transported in the dark back to the lab (< 2 h) then 100 mL was filtered through a 25 mm glass fiber filter (GF/F; 0.7 um) and stored in the dark at −20°C until processing. Chlorophyll-a was extracted with 90% acetone and quantified by fluorometry ^31^.

### PlanktoScope imaging and analysis

We built and replicated a PlanktoScope v2.1 as previously described ^26^. The 2021 fieldwork served as a trial deployment since the standardized protocol was not yet available. In 2022, imaging was done following the standardized protocol v1^32^ wherein the pump volume and image size were calibrated. Hence, only the 2022 counts are reported, but both datasets provided insights into morphological features that can discriminate plankton groups. On average, we collected 1000 images per sample, totaling ~42,900 images covering 23 sampling trips. The machine records the total imaged volume used for cell count estimates. We labeled 1250 representative images as the training set. Plankton identification was based on comparing cell gross morphology to images in existing identification guides ^33,34^ with an emphasis on discriminating total diatom vs. total dinoflagellates which were the most distinctive groups and the groups responsible for HABs in this region. We implement image contrast normalization to ensure backgrounds with consistent color. We trained a Residual U-Net ^35^ to perform semantic segmentation on the labeled data using pixel-level BCELoss, followed by noise filtering, contour-finding algorithm ^36^ and entity counting. We performed the counting using the trained model seven times to get diatom and dinoflagellate aggregate estimates.

### Phytoplankton microscopy counting

Microscopy counting was performed in parallel with PlanktoScope counting on eight select samples (four from station 1, four from station 4) during the April-May 2022 season. Additional microscopy counts were conducted on a different set of samples collected biweekly at Station 1_CCMS (Feb-Apr 2022; Jan-Oct 2023). For microscopy, a 1-mL aliquot of sample was deposited onto a Sedgewick Rafter slide and cell counts were performed thrice per sample bottle and then averaged. Identification was made down to the genus level based on existing guides ^33,34^.

### Electron microscopy of Bolinao VLPs

Viruses in 2 mL of the 200 µm pre-filtered seawater were concentrated using Amicon Ultra-0.5 30kDa (12/04/2021) and 16 mL in Amicon Ultra-4 100kDa (06/06/2022) centrifugal ultrafilters (Millipore Sigma). The concentrated samples were preserved in 2% glutaraldehyde at −20°C until processing. Grids (200 mesh copper with carbon-stabilized formvar) were rendered hydrophilic by glow discharge. A 4 µL sample was applied to the grid and viruses allowed to adsorb for 45 seconds. Sample was then wicked away with filter paper followed by washing with distilled water and negative staining with 2% uranyl acetate. The grid was viewed on a Hitachi HT7700 TEM at 100 kV and photographed with an AMT XR-41B 2k x 2k CCD camera.

### CCMS-AWQMS data collection

The Automated Water Quality Monitoring Station (AWQMS) at Station 1 is equipped with sensors for conductivity, temperature, and depth (CTD), and DO. In situ measurements were recorded hourly. Temperature, salinity, and chlorophyll were measured by sensors that sampled seawater through an inflow tubing, while the DO sensor was directly submerged. These measurements complemented the data collected using the handheld multiparameter probe noted above.

### 18S rRNA gene amplicon analysis

DNA was extracted from the 0.2 µm polycarbonate filters using a MoBio PowerSoil DNA Isolation kit. The V9 region of the 18S rRNA gene was amplified with the Earth Microbiome Project primers (1391f-GTACACACCGCCCGTC and EukBr-TGATCCTTCTGCAGGTTCACCTAC) then sequenced on the Illumina MiSeq platform to generate 300 bp paired-end (PE) reads ^37,38^. Sequencing was done at the University of Hawaii Advanced Studies in Genomics, Proteomics and Bioinformatics Sequencing Facility. Using the software package Mothur v1.48 ^39^, PE reads were assembled, quality filtered, and aligned to the SILVA database v132 ^40^. Sequences were checked for chimeras and clustered into operational taxonomic units (OTUs) at a 97% similarity. Sequences were taxonomically classified using SILVA v132. Community comparisons and statistical analysis were implemented in Mothur using subsampled libraries to keep the number of sequences the same across samples. Additional analyses at the level of amplicon sequence variants (ASVs) or phylotype (genus) were done. Non-metric multidimensional scaling (NMDS) plots illustrating the relationships among samples were made using the vegan package in R.

### Metagenome preparation

DNA was extracted from 0.02 µm Anotops using a Zymo DNA Microprep kit, but modifying the protocol to include proteinase K and a backflushing of the lysis buffer through the filter in the housing^41^. Lysis buffer recovered from the filter housing with a syringe was deposited into a microtube and the remainder of the purification steps followed the manufacturer’s protocol. The resulting DNA samples were sent to Azenta, NJ, USA, for 150 bp PE Illumina HiSeq metagenomic sequencing. Raw reads were subjected to adapter trimming and quality filtering using Trimmomatic ^42^. Reads were normalized using bbnorm (minimum 5x and average depth of 50x) (https://sourceforge.net/p/bbmap). A mixed co-assembly approach was implemented using MEGAHIT ^43^. Reads in the same station and month were combined for co-assembly (**Table S1 & S2**). All the resulting assemblies were combined in one file, and contigs less than 1000 bp were removed using SeqKit ^44^. Contigs were clustered at 95% nucleotide similarity using CD-HIT ^45^ to remove the redundancy, resulting in 2.28 million contigs. Before the downstream analysis, the assembly file was re-formatted to simplify names using Anvio ^46^. 28,462 bins/MAGs were produced through Bowtie2 mapping ^47^ and MetaBAT2 binning ^48^. DAS tool was used to identify the contigs-to-bin memberships ^49^ and SeqKit was used to determine the length statistics ^44^.

### NCLDV identification, characterization, and co-occurrence network

Viral contigs were identified using geNomad ^50^ producing 1,739 candidate NCLDV bins. Only those bins with sizes between 100kb - 3Mbp were retained, corresponding to the known smallest (103 kbp ^51^) and largest (2.77 Mbp ^52,53^) NCLDVs. We retained only those bins with < 40 contigs, similar to a previous study ^54^. We also included only those bins with the longest contig >10,000 bp. These length cutoffs resulted in 190 bins with a lower degree of fragmentation. ViralRecall ^55^ was used to identify the five highly conserved NCLDV marker genes, including MCP (major capsid protein), polB (DNA polymerase B), A32 (A32-like packaging ATPase), VLTF3 (virus-like late transcription factor), and SFII (superfamily II helicase). Only the bins with 4 out of 5 marker genes were retained, resulting in 90 bins. Additional NCLDV marker genes were identified, including RNAPL, RPANS (RNA polymerase large and small subunits, mRNAc (mRNA capping enzyme), RNR (ribonucleotide reductase) and D5 (D5 primase/helicase). Only those contigs with positive ViralRecall scores were retained in each bin. In addition, we removed bins with greater than 7 MCP copies to lessen the chance of analyzing chimeric bins. It is reported that NCLDV can contain multiple MCP copies, for instance, BpV1 has 5 (NCBI: NC_014765.1), PBCV-1 has 7 (NC_000852.5), OtV5 has 8 (NC_010191.2). These criteria led to a final set of 82 bins. Genome completeness is estimated using CheckV ^56^ and rRNA genes were identified using barrnap (https://github.com/tseemann/barrnap). No rRNA genes were found in the final set of 82 GV-MAGs. Genome annotation was done using NuMP (https://github.com/BenMinch/NuMP; accessed Jan 2024) and ViralRecall ^55^. KEGG metabolic module completeness was calculated using MicrobeAnnotator via DIAMOND blastp ^57^. MCP and PolB phylogenetic trees were constructed using NuPhylo (https://github.com/BenMinch/NuPhylo; accessed Jan 2024) and were visualized in iTOL ^58^. Taxonomic identification was done using the phylogenetic trees and confirmed using TIGTOG ^59^. Anvio was used to calculate the mean coverage^46^. The normalized relative abundance of bins is calculated by dividing the number of recruited reads by the total number of reads. The number of recruited reads is calculated by multiplying Anvio’s mean coverage by bin length divided by read length. Bakta was used for genome visualization ^60^. Finally, after removing diatom and zooplankton 18S OTUs, a co-occurrence network between remaining phytoplankton and GV MAGs was made using SpiecEasi in R ^61^. The resulting network was visualized using igraph and ggplot.

## Results

### Environmental condition, saxitoxin episode, and fish kill

Intensive mariculture activities within the Guiguiwanen Channel in Bolinao **(Figures 1 and S1)** creates a DO and pH gradient over a short distance from the fish cages to the channel mouth **(Figures 2 and S2)**. Our sampling covers different seasons and in 2021, the Philippine weather agency reported the onset of the NE monsoon on October 25 ^62^ with a surge reported on December 2, leading to a decrease in surface seawater temperature **(Figure 2)** ^63^. Otherwise, surface temperature, salinity, and TDS were similar across sites and seasons **(Figures 2 and S2)**, except at the start of the saxitoxin (STX) episode (April 13, 2021), when a drop in salinity to ~26 PSU was observed. Chlorophyll-a, a rough proxy for phytoplankton biomass, varied from 1–20 ug L^−1^ but was relatively higher during NE monsoon **(Figures 2 and S2)**.

**Figure 2.**
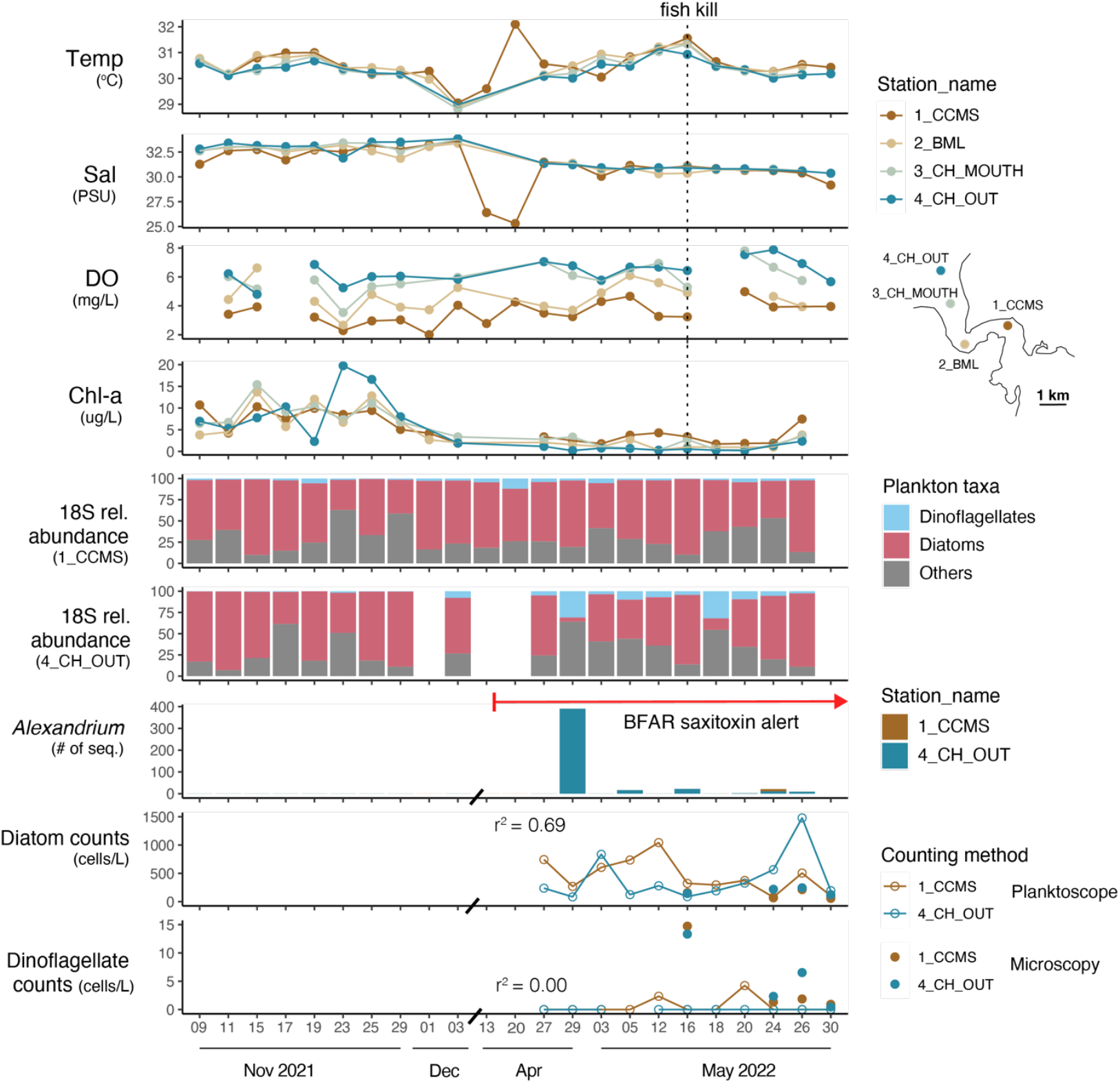
Time series of physical, chemical, and phytoplankton data. [From top to bottom] Plots showing similar surface temperature and salinity ranges. A decrease in salinity was also observed at the onset of saxitoxin alert. In contrast, a gradient of dissolved oxygen was observed. Chlorophyll-a fluctuates, reaching up to 20ug/L, and is relatively higher in Nov-Dec 2021. Diatoms are the most abundant phytoplankton; however, in some cases, dinoflagellates reach relatively high abundance (Station 4; April 29 & May 18). *Alexandrium* appearance coincides with BFAR saxitoxin alert. No *Pyrodinium* sequences were detected, while *Gymnodinium* was present and in low 18S quantity. Finally, PlanktoScope counts agree with microscopy counting for diatoms (r^2^=0.69) but had difficulty counting dinoflagellates accurately because of their relatively small size. Moreover, a fish kill event happened on May 15-16, 2022.

During the second sampling, there was a 2.5-month period of paralytic shellfish poisoning (PSP) alert triggered by elevated STX levels in shellfish tissues. Persistent detection of STX in the shellfish started on April 7, 2022. The toxin level was reported at 249 µg STX per 100 g shellfish at the start of the ban, beyond the 60 µg STX per 100 g shellfish limit, as reported by the local monitoring agency (BFAR) ^64^. The shellfish bulletin (herein referred to as the BFAR toxin alert) was only cleared on June 17, 2022. A fish kill event was recorded on May 15, 2022, with images from May 16 showing extensive fish mortality near Station 1 **(Figure S3)**. Surface DO at Station 1 remained consistently low for months, sometimes reaching hypoxia (<2 mg/L) (**Figures 2 and S4**).

### Phytoplankton monitoring and composition

Phytoplankton monitoring was conducted using three methods: [1] 18S rRNA amplicon sequencing, [2] a PlanktoScope **(Figure S5)**, and [3] conventional microscopy identification and counting. The 18S rRNA sequencing provided relative abundances of dominant phytoplankton with high taxonomic resolution, while PlanktoScope provided cell count estimates with relatively low taxonomic resolution (**Figure S6)**. Our analysis focused on Stations 1 and 4 because of logistical constraints. The PlanktoScope captured images of dominant plankton and our taxa of interest—dinoflagellates and diatoms.

### Plankton diversity and succession revealed by 18S rRNA gene sequencing

Diatoms frequently dominated the amplicon pool, comprising 5–97% (66% on average) of 18S rRNA gene sequences over the sampling period. However, on April 29 and May 5, 2022, dinoflagellate amplicons (31% and 32% of the sequences) were more abundant than those of diatoms at Station 4 (**Figure 2 and S7)**. Phytoplankton community composition significantly varied by season and, to some degree, station **(Figure 3A)**, with consequent influences on giant virus composition **(Figure 3B)** as discussed in the next section. Species richness and diversity increased from the NE monsoon to SW monsoon **(Figure 3C)**. Specifically, we observed a conspicuous diatom succession from *Skeletonema* (early November), *Chaetoceros* (late November), unclassified Mediophyceaea (end of November), *Thalassiosira* (December) back to *Skeletonema* (April), then unclassified Diatomea (early May), and mostly *Chaetoceros* (end of May) **(Figure S8)**. Succession in the dinoflagellate community was difficult to resolve as most sequences belonged to unclassified Dinophyceae and Gymnodiniphycidae **(Figure S9)**. Still, several dinoflagellate genera were evident, such as *Gyrodinium, Karlodinium, Scrippsiella, Heterocapsa, Protoperidinium*, and *Alexandrium* **(Figure S9)**. Other notable plankton included Chlorophyta (0.29-34%) represented by *Ostreococcus, Micromonas*, and *Tetraselmis* **(Figure S7)**. Non-phytoplankton groups that were captured by 18S rRNA gene sequencing include copepods (0-35%), tunicates (0-2.8%), and ciliates (0.14-11.6%) **(Figure S7)**.

**Figure 3.**
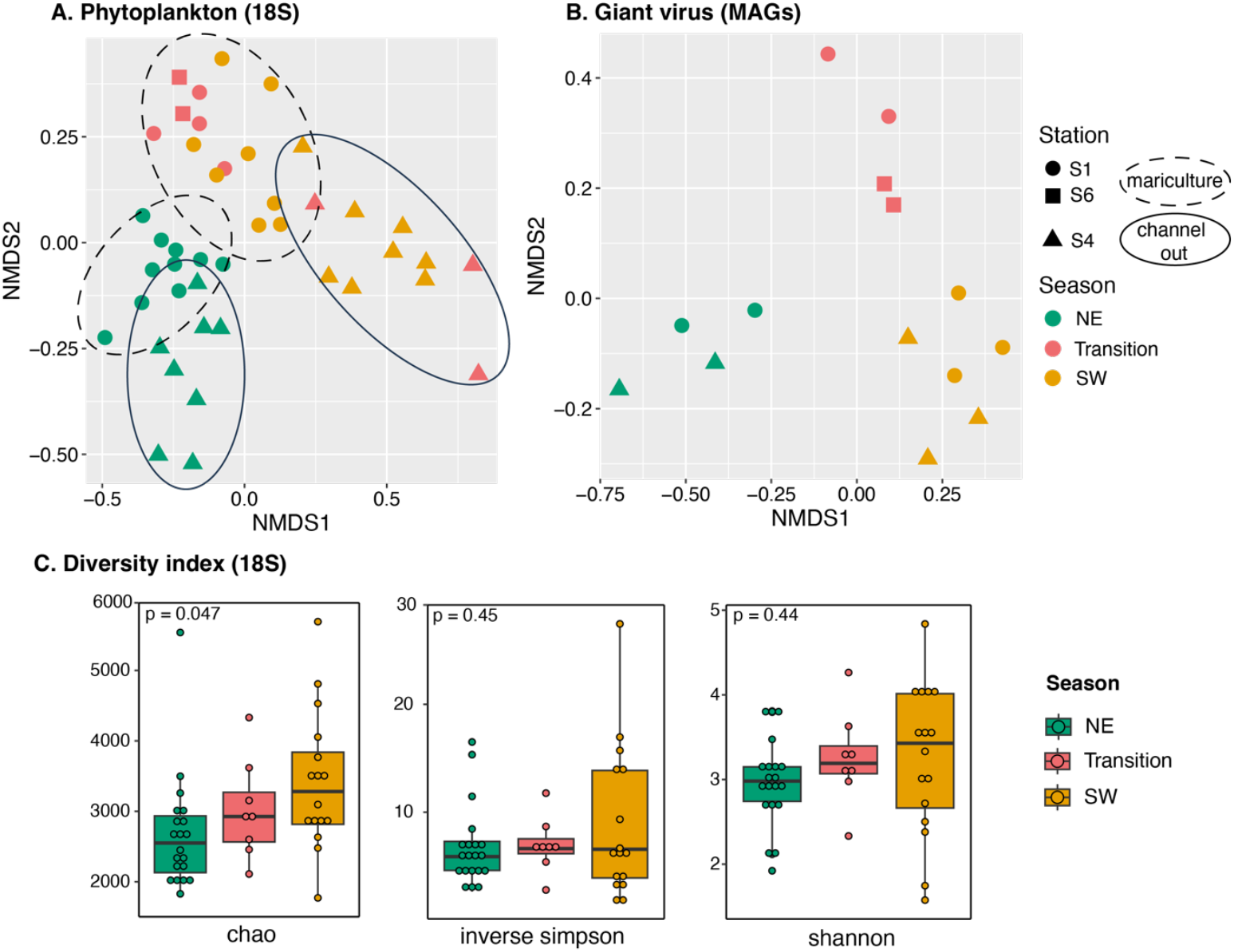
Community structure and diversity of phytoplankton and giant viruses. Non-metric multidimensional scaling plots illustrate seasonal and spatial variations in (A) phytoplankton communities based on 18S rRNA gene amplicons and (B) giant viruses based on metagenome-assembled genomes (GV-MAGs). Symbols are colored according to season and symbol shapes reflect the stations from which samples were collected. (C) The richness and diversity indices of phytoplankton derived from 18 S rRNA gene amplicon data show an increasing trend from NE monsoon, though the transitional period, to the SW monsoon. P-value based on ANOVA test.

*Alexandrium* sequences constituted a small fraction of the plankton community (at most 0.1% of 18S rRNA gene sequences). Of the three STX-producing genera, *Alexandrium, Pyrodinium*, and *Gymnodinium* ^65^, only *Alexandrium* coincides well with the BFAR STX alert **(Figure S10)**. *Pyrodinium* was absent throughout the time series. Further analysis at the OTU and ASV levels suggested the presence of multiple *Alexandrium* represented by two OTUs (Otu00725 & Otu01700) and seven *Alexandrium* ASVs (two are singletons not included in the figure) **(Figure S11)**. Microscopy data supported these findings, showing that *Alexandrium* sp. was present at 83–111 cells/L during the start of the shellfish ban but represented only 0.15–0.38% of the total phytoplankton community (**Figure S12)**.

### PlanktoScope performance

The correspondence of PlanktoScope and microscopy counts is better for diatoms (linear regression, r^2^ = 0.69, n=8) than the small dinoflagellates (<40 µm) **(Figure 2)**. This is likely due to the limits of PlanktoScope optical resolution which is best suited for plankton in the 40–200 µm range ^32^. Nevertheless, we can still detect the essential features of dinoflagellates, such as girdles and horns, but not the thecal plates **(Figure S6)**. Microscopy identified 33 plankton taxa during the 2022 sampling **(Table S3)**. Station 1 had seven dinoflagellate genera, eleven diatom genera, and four zooplankton taxa, with diatoms—especially *Pseudo-nitzschia, Skeletonema, and Thalassiosira*—being the most abundant. Station 4 had ten dinoflagellate genera, twenty-five diatom genera, one cyanobacterium, and six zooplankton taxa, with *Pseudo-nitzschia* also dominating, followed by *Chaetoceros* and *Skeletonema*. Other notable forms in both stations were the dinoflagellate *Peridinium* sp. and zooplankton forms, bivalve larvae, and tintinnids.

### Diversity and dynamics of Bolinao giant viruses

Giant viruses (GVs) were investigated using metagenomics to examine their diversity and potential interactions with phytoplankton. Based on our filtering criteria, which includes genome size and presence of NCLDV markers, we recovered 82 taxonomically diverse GV MAGs spanning five orders: *Algavirales, Imitervirales, Pimascovirales, Pandoravirales*, and the recently suggested new class/order *Proculvirales*/-*viricetes* (PC_01) **(Figures 4A, S13, S14; Table S4)**. The GV MAGs were further classified into 13 families: AG_01 (Prasinoviruses), AG_04, IM_01 (*Mesomimiviridae*), IM_09 (*Schizomimiviridae*), IM_12 (*Allomimiviridae*), IM_13, IM_16 (*Mimiviridae*), PM_01, PM_06, PM_07 (*Pithoviridae*), PV_03 and PV_05 (*Coccolithoviridae*) **(Table S4)**. One GV MAG (GV78) was putatively classified within newly identified taxa, PC_01, thought to be exclusive in the Southern and the Arctic Oceans ^66^. These GV MAGs have varying CheckV quality and completeness from high quality (n=18) and medium quality (n=41) to low quality (n=24) **(Table S4)**, with the largest (GV18) being 960 kb and smallest (GV82) with a 101 kb genome. Electron microscopy confirmed the presence of virus-like particles (VLPs), with diverse morphologies including ovoid, filamentous, isometric, spindled, and tailed VLPs **(Figure S15)** and large icosahedral giant virus-like particles (~170-200 nm) **(Figure 4B)**.

**Figure 4.**
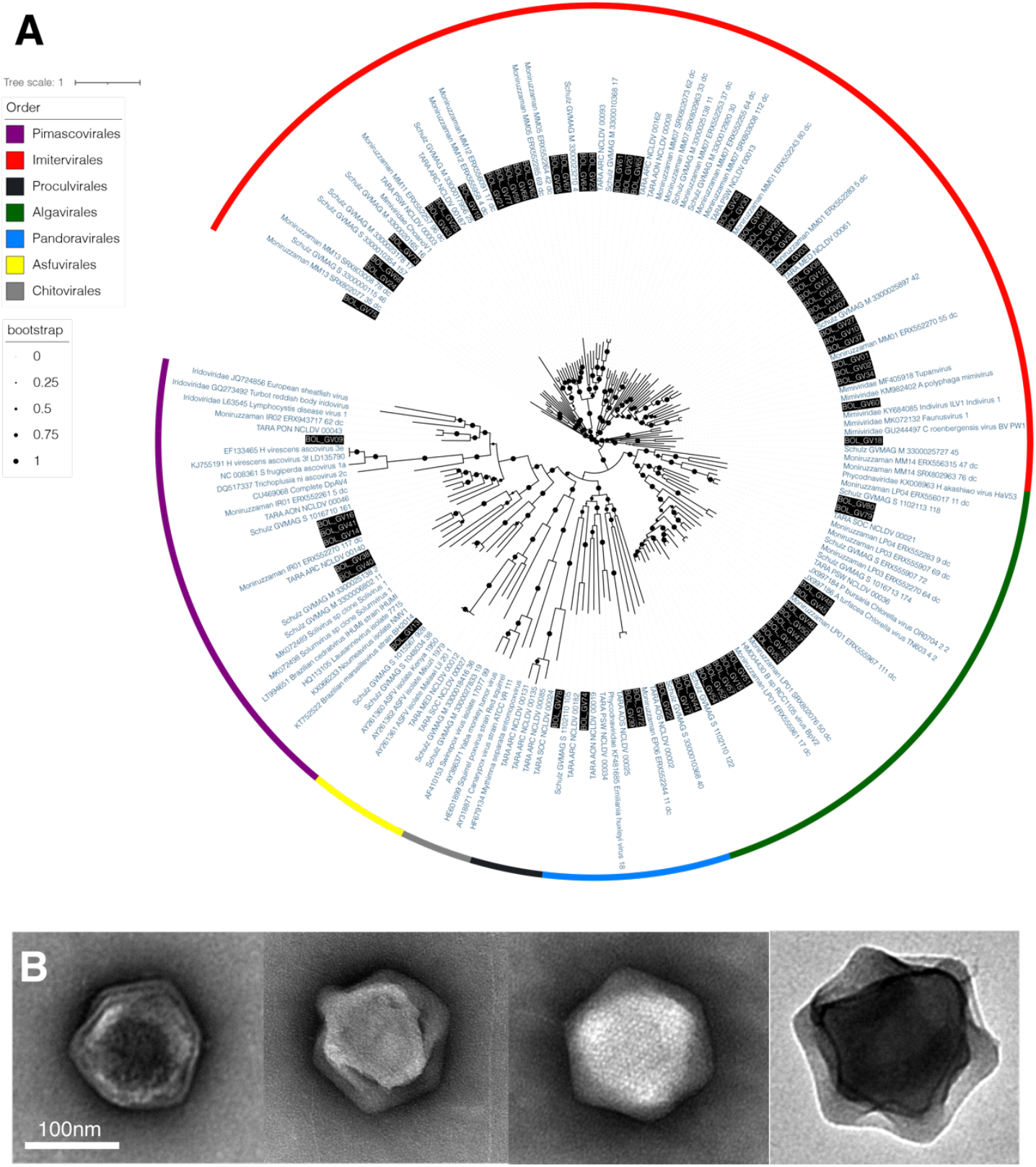
Phylogeny and morphology of Bolinao giant viruses. **(A)** Phylogeny of GV MAGs based on PolB sequences. Highlighted in black are the Bolinao GV MAGs. Written in dark blue are representative sequences from different NCDLV families, as provided by the NuPhylo package. For complete taxonomy, see supplementary materials. **(B)** A survey of viral-like particles (VLPs) in Bolinao waters shows VLPs with sizes ~170-200 nm.

These GV-MAGs exhibit extensive metabolic capacities, with genes involved in glycolysis, gluconeogenesis, tricarboxylic acid (TCA) cycle, light harvesting/photosynthesis, cytoskeleton, transporter, and nutrient metabolism **(Figure 5)**. One GV-MAG code for an ammonium transporter (GV17) and six coded for phosphate transporters (GV11, GV19, GV21, GV28, GV57, and GV71). **(Figure 6A)**. The relative abundance of GV-MAGs varied seasonally, with three main clusters, representing roughly the months of sampling and monsoon season—November (Cluster 1; NE monsoon), April (Cluster 2; Transition period), and May (Cluster 3; onset of SW monsoon) **(Figure 6A)**. We further examined the functional capacity of our GV-MAGs by estimating the presence and completeness of KEGG metabolic modules **(Figure S16)**. Present in the transition period and SW monsoon clusters are the beta-lactam biosynthesis pathway, cysteine and methionine metabolism and drug resistance pathways **(Figure S16**). Among all the GV-MAGs, the largest (GV18) had the most extensive metabolic capacity and the most copies of glycolysis and pentose phosphate pathway genes **(Figure S17)**. GV01 and GV02 were notable for containing several copies of ‘chitin response’ and lipid metabolism genes. GV01 contains a rhodopsin and a chlorophyll AB binding protein, while GV02 contains carbonate anhydrase. GV02 also encodes for a sugar transporter (MtN3_slv) and apoptosis inhibition-associated gene (Bax1-I).

**Figure 5.**
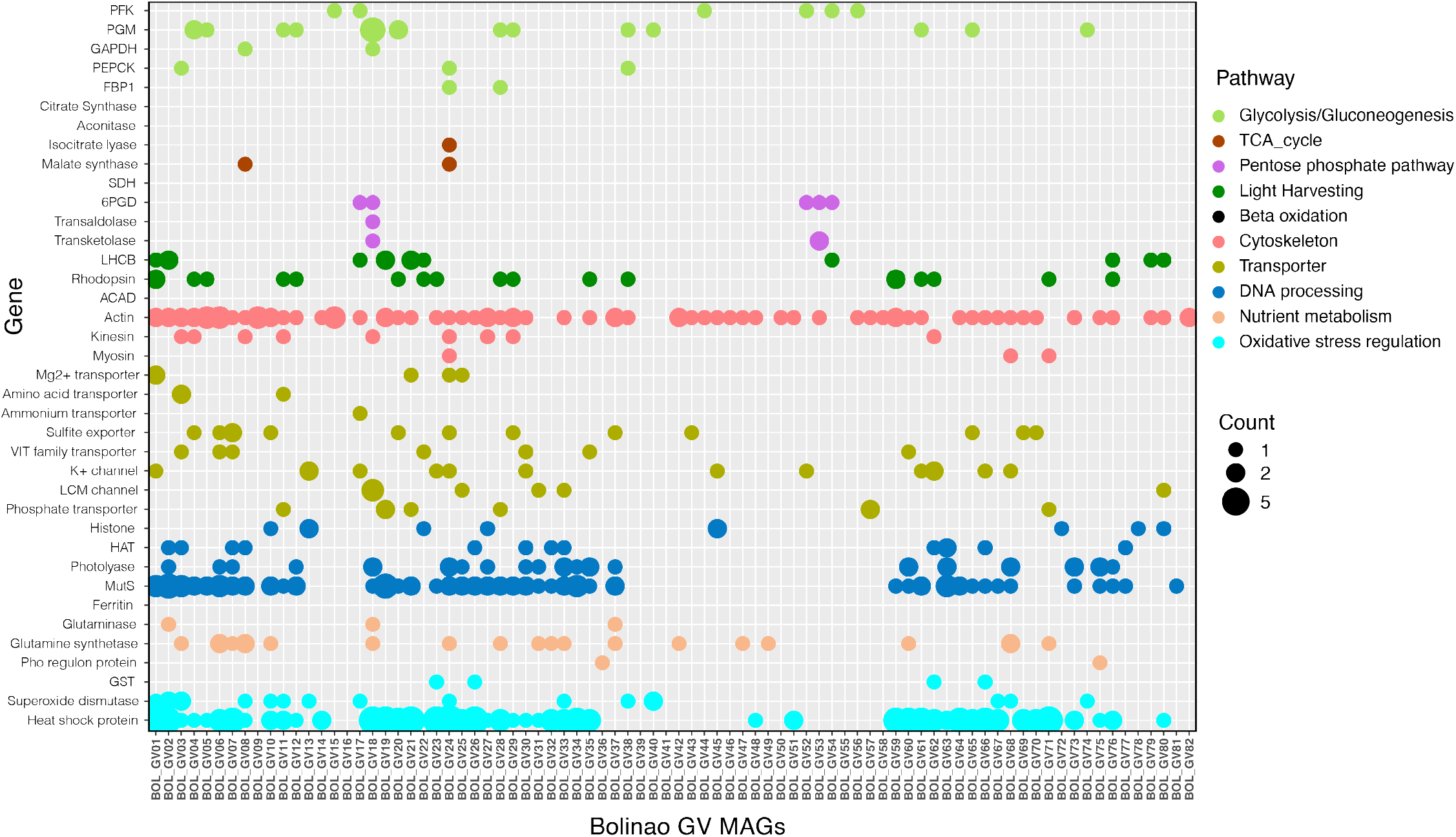
Auxiliary metabolic genes and virologs of Bolinao GV MAGs. Curated auxiliary metabolic genes or virologs (colored based on pathways). The bubble size is based on the number of copies in the genome. The gene set is based on Ha et al. 2021 and implemented in the NuMP package.

**Figure 6.**
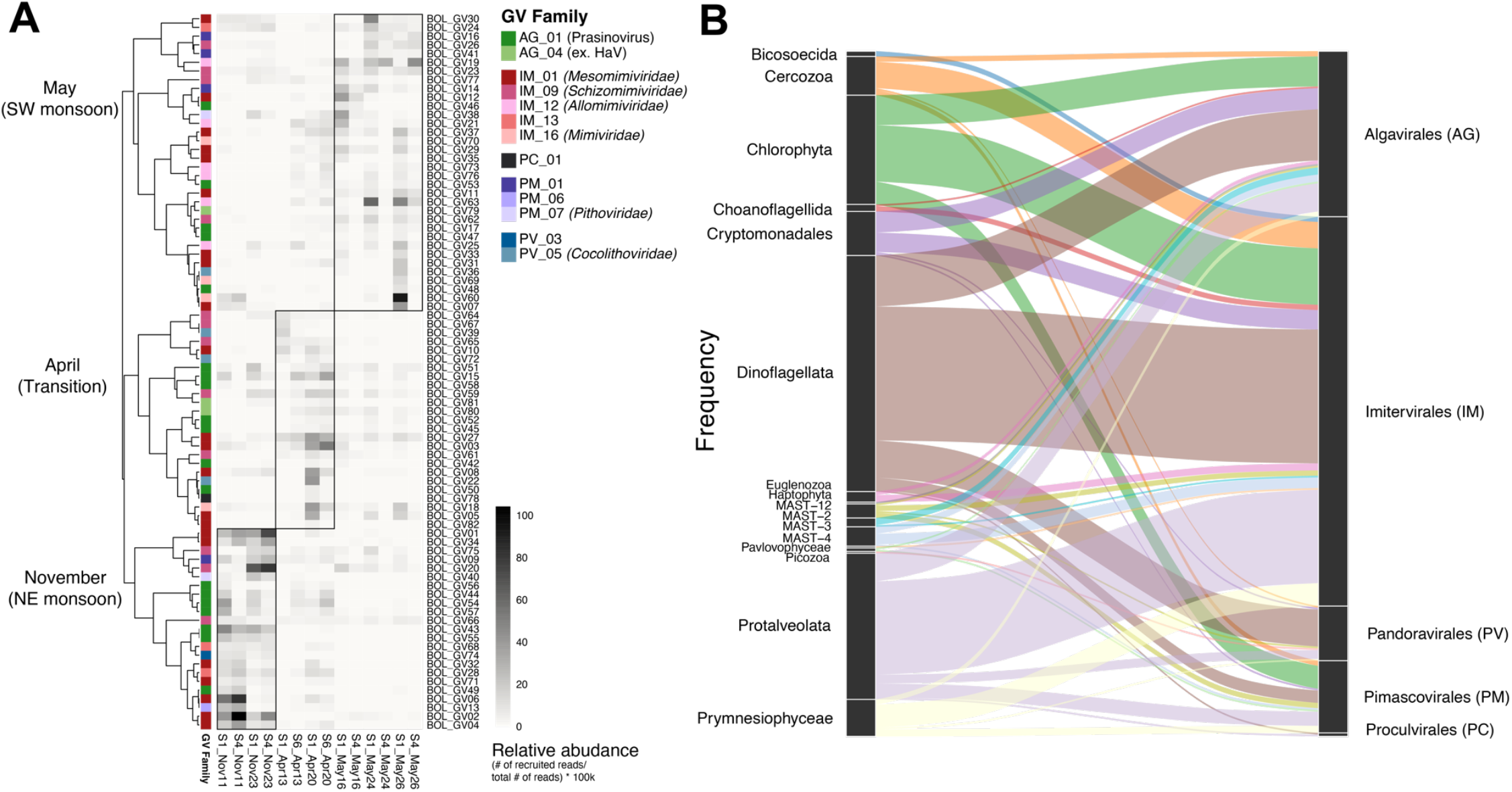
Temporal GV dynamics and phytoplankton-GV associations. (A) Heatmap showing the relative distribution of GV MAGs and (B) Frequency of plankton-GV associations based on their corresponding taxa.

We analyzed the co-occurrence of phytoplankton and giant viruses to gain insight into putative host-virus associations linking 495 phytoplankton OTUs with 82 giant viruses **(Figure 6B and S18; Table S5)**. We focused on phytoplankton hosts known to be infected by giant viruses; thus, we removed diatom and zooplankton OTUs. Known virus-host pairs are captured in the network. The high frequency of green algae and dinoflagellate co-occurring with *Algavirales* and *Imitervirales* (known viral orders infecting algae) is of interest. For instance, *Ostreococcus* (Otu00039) co-occurs with GV53, family AG_01, the known Prasinovirus family. Another example is Prymnesiophyceae (*Chrysochromulina*; Otu00104 & Otu01410), which co-occurs with IM_01 (in which Chrysochromulina parva virus belongs). One *Tetraselmis* (Otu000169) co-occurs with three Algavirales GV15, GV54, and GV51. The relative abundance of *Alexandrium* was too low to be captured by the network, but several dinoflagellate OTUs, such as *Gyrodinium* (Otu00034), *Karlodinium* (Otu00209), *Pelagodinium* (Otu00906), and *Heterocapsa* (Otu00213), were found to be associated with GVs belonging to *Imitervirales* families.

## Discussion

The Bolinao mariculture area has experienced repeated eutrophication-induced HABs for years ^3,9^ as a result of the high concentrations of fish structures and nutrient input from excess fish feed. Consequently, DO and pH vary over a short distance from mariculture-intensive area to the channel mouth likely due to the decomposition of organic matter that consumes oxygen and produces carbon dioxide that lowers pH ^67^. During this study, the causative toxic HAB species was likely *Alexandrium* sp., as supported by our observations (~0.1% 18S sequences, ~0.35% phytoplankton counts, around 83–111 cells L^−1^) and consistent with historical accounts ^5,7,68^. This is in contrast to many other areas of the Philippines (e.g. Sorsogon Bay, and Eastern Visayas embayments, and Dumanquillas Bay to name a few) where *Pyrodinium* is the primary STX-producing species ^69–72^. Even low concentrations of toxin-producing cells can result in high risks of PSP. In a 2014 PSP episode involving *Alexandrium tamiyavanichii*, cells reached only 840 cells L^−1^ and resulted in 3,500 µg STX per 100 g shellfish in Kuantan port, Malaysia ^73^. Likewise, in Bolinao, *Alexandrium minutum* concentrations of only 5–7 cells L^−1^ were associated with accumulation of 930–2,894 µg STX per 100 g shellfish meat ^74^. This is consistent with our observation that, despite a PSP alert, *Alexandrium* was a minor component of the phytoplankton community at the time. This highlights the need for rapid, sensitive, and specific methods for detecting toxin-producing species.

Typically, a drop in salinity and DO is observed during HABs and fish kill in Bolinao. The 2010 *Alexandrium* toxin bloom was associated with low salinity (26–27) and low DO (<2.8 mg L^−1^, as low as 0.3 mg L^−1^ at the bottom) ^8^, consistent with the current study. *Alexandrium* species are known to survive and produce more toxins at low salinity ^75^. The observed DO decline during this study was also associated with a fish kill. These DO estimates are surface measurements; thus, we expect an even lower DO at the deeper portion of the water column, which could be lethal for fish, especially demersal species. DO declined several times in a 2-year period, but not every drop in DO leads to a fish kill (**Figure S4**); thus, the May 15 fish kill event may have other, or at least additional, causes such as toxins, ammonia poisoning, or gill obstruction.

The overall phytoplankton community observed in this study is typical of eutrophic coastal areas dominated by diatoms and dinoflagellates ^76,77^ along with other common marine taxa such as chlorophytes and haptophytes ^78^. A 2018 Bolinao survey reported similar composition, with dinoflagellates (~35–65%) and diatoms (~30–60%) as the dominant taxa in the inner Guiguiwanen channel ^27^. Monsoon seasons were the major driver of community changes consistent with previous work ^79,80^. The relative uniformity across stations within a season may reflect the close spacing between the sampling sites (a few km).

Aside from the 18S amplicon approach, we tested the utility of PlanktoScope, a community-driven, open-access, low-cost camera ^26^. We illustrated the applicability of the machine for estimating diatoms but had less success counting and identifying dinoflagellates. Increasing the optical resolution, and perhaps refining the detection algorithm, could improve this situation. A newer, more stable version of the PlanktoScope machine (v2.6) became available after completion of this study (we used v2.1). Adoption of this newer model might also improve count accuracy. The PlanktoScope, being the most economical and accessible machine (compared to FlowCam and Imaging FlowCytobot), might be a means to improve monitoring coverage, especially in resource-poor settings. The one described in this study represents our first attempt at utilizing PlanktoScope in monitoring HABs and has shown promise.

In addition to phytoplankton, we examined GV diversity and its spatiotemporal variation. As expected, the Bolinao GV MAGs encoded extensive suites of metabolic genes, including nutrient transport that could transiently alter the nitrogen and phosphorus acquisition capabilities of the infected host cell (the virocell). Such a virus-mediated alteration of host phenotype was observed during infections of the green alga *Ostreococcus tauri* with the virus OtV6. In that study, the virus-encoded ammonium transporter expressed during infection appears to have a higher affinity, but lower V_max_, for methyl-ammonium than the host-encoded transporter, which is presumed to be advantageous in nitrogen-limited waters ^81^. The Bolinao GV MAGs encoding for ammonium (n=1) and phosphate transporters (n=6), may similarly reflect adaptations to nitrogen or phosphate limitation in Bolinao waters ^3,9^. The higher frequency of phosphate transporters may be a reflection of the high nucleic acid content of virions, which increases their relative P requirement compared to that of the cells in which they replicate^82^. The Bolinao GV MAGs also encode a whole suite of energy-generating metabolic enzymes related to photosynthesis, glycolysis, pentose phosphate pathway, and TCA cycle, which presumably either augment or replace host functions during energy-demanding processes of viral replication. One GV MAG encodes for carbonic anhydrase that can influence the carbon concentrating mechanism and perhaps aid photosynthesis in the phytoplankton virocell ^83,84^. In the clusters representing the transition period and SW monsoon samples, KEGG pathways related to antioxidants are enriched, which may reflect selection for viruses that have capacity to counter oxidative stress during infection ^85,86^. In addition, KEGG pathways related to multidrug resistance efflux pumps are also enriched during these months, which might be related to removal of antiviral toxins ^87^.

The giant viruses described here were likely not involved in direct bloom termination, as the phytoplankton community was dominated by diatoms which seem to be infected only by small viruses ^88^. However, the putative phytoplankton-GV associations described here, and their changes over time, suggest that the GV are involved in community dynamics during Bolinao HABs, similar to what is observed in HABs elsewhere ^89^. GV could be involved in the demise and succession of HABs at other times, especially for blooms dominated by dinoflagellates for which some giant viruses are known. In such cases, viruses might be more efficient in killing HAB species than zooplankton which are usually deterred by toxins ^90,91^. Several single species blooms have occurred in Bolinao. For instance, in 2002, a mono-species bloom of *P. minimum* (99%) lasted several days, succeeded by diatoms (87%) after 4 days. In 2010, an *Alexandrium* sp. bloom (~86%) succeeded by a *S. costatum* bloom (99%) was also recorded. In 2018, a mono-species *Takayama* sp. bloom was associated with a fish kill event ^27^. Aside from the direct impact on the bloom, viral infection will have pronounced consequences on carbon export and microbial succession. For instance, it was shown that viral infection causes a 2-4-fold increase in carbon release and a rise of not only bacteria but also eukaryotic heterotrophic organic matter recyclers ^92^.

By focusing only on large dsDNA-containing viruses, we did not account for other viruses that may have influenced bloom dynamics, especially those infecting diatoms. Most diatom viruses are small and contain ssRNA or ssDNA ^88,93,94^. Nonetheless, this study expands our knowledge on giant virus diversity and spatiotemporal dynamics in the context of an STX episode and across a eutrophic coastal environment and monsoonal seasons. Some research groups are exploring viruses as HAB mitigation.

One study has shown a significant reduction in the density of the harmful algae, *H. circularisquama*, with the application of either virus alone or virus-sediment mix in a mesocosm experiment ^95^. This underscores the urgency and importance of understanding phytoplankton-virus interaction and dynamics in various contexts to fully comprehend the impact of such mitigation efforts.

## Conclusion

This study examines phytoplankton diversity and identifies associated giant viruses during different monsoon seasons, a toxic bloom, and a fish kill, contributing to the growing literature on HAB dynamics. We used a complementary method for phytoplankton/HABs tracking that provided data on *Alexandrium* sp. appearance coinciding with STX accumulation in shellfish. Further fine-tuning of PlanktoScope to better detect and count dinoflagellates in addition to diatoms will increase the efficiency of HAB monitoring. We documented the diversity of giant viruses, but further work is needed to link specific viruses and their hosts and determine the influence of viral lysis on the bloom dynamics. Bolinao HABs remains a recurring problem and detailed investigations into the various sources of algal mortality may shed light on the variables influencing bloom severity and duration.

## Acknowledgments

This project was funded by the National Geographic Early Career Grant (EC-KOR-66418R-21), East-West Center Field Research Grant, Experiment’s Ocean Solutions Grant, and Uehiro Fellowship to APG. We are indebted to Tina Carvalho at the Microscopy Facility at the PBRC for all the electron microscopy training and usage. We thank the Rappé lab (UHM) and Dr. Charissa Ferrera (UPD) for providing some materials and lending equipment. We are thankful for the Microbial Oceanography Lab, led by Dr. Deo Onda (UPD) through the UPGRADE CIA project funded by the National Security Council for providing supplies and a DNA extraction kit. The PlanktoScope was made with the help of Jie Zhou (Fabrication Lab, UHM College of Engineering), Ian Quiño Fernandez, Eric Kolb, Dr. Manu Prakash, Dr. Adam Larson, Thibaut Pollina, and the rest of the PlanktoScope community. Niño Posadas, Jake Baquiran, Dr. Malou San Diego-McGlone, Miraflor Sanchez, Dr. Josefa Pante, Dr. Van Rodriguez, and Abby Melendres were pivotal in completing the required paperwork, research agreements, and permits. Erika Gernato and Raffi Isah assisted in obtaining materials, reagents, and instruments. We are also grateful to Cha Caalim, Rubie Esmolo, Angel Doctor, and the rest of the Bolinao Marine Lab research assistants and boatman for the marine lab access, sampling, and conducting community outreach in Bolinao.

## Data Accessibility

The data used in this paper have been deposited in the NCBI Short Read Archive as BioProject PRJNA1237401 for 18S sequences and BioProject PRJNA1238504 for metagenomic sequences. The GV-MAGs assembly and annotation are available at FigShare https://figshare.com/s/79c4bebb28187961279b. The codes for PlanktoScope counting are available at https://github.com/jjshoots/PlanktonSegmentationTrainer.

## Author Contributions

APG, CC, ATY, and GFS conceived of the study. APG and GCMBA performed the sampling and experiments. APG, GCMBA, JJT, and GAB performed the analysis. ELDG, RBF, CLAR, and CLV contributed data. APG and GFS wrote the paper. All authors edited and approved the manuscript.

