## Supplemental Material 1 for "Phytoplankton and giant virus dynamics during different monsoon seasons, a fish kill, and a toxic bloom in a eutrophic mariculture area"

### Supplementary Materials

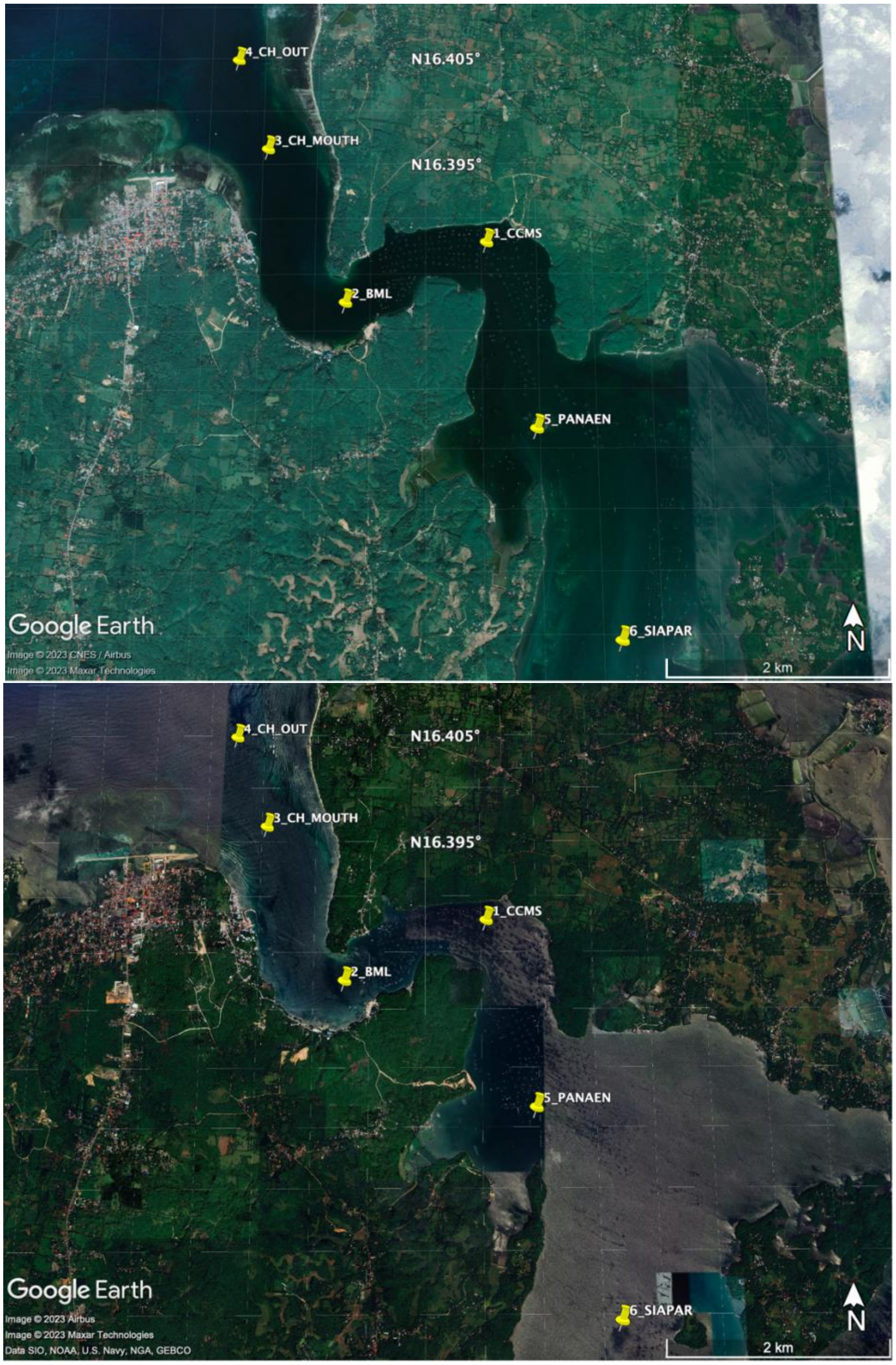


**Figure S1. Study sites in Nov 2017 (top) and Dec 2023 (bottom) showing the location of the fish structures.** In these images, the fish structures are shown as circular or rectangular pens located inward of the channel from Station 2 BML to Station 6 Siapar.

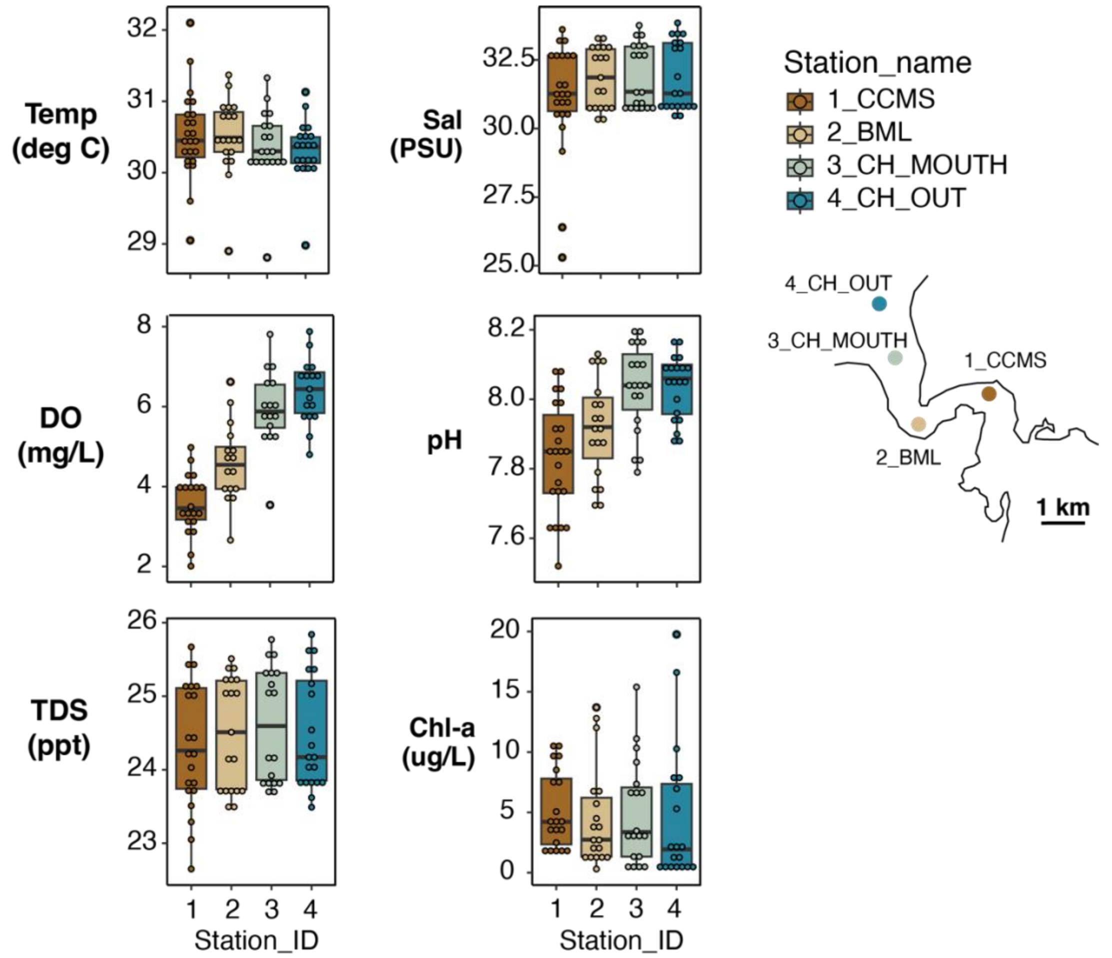


**Figure S2. Bolinao physicochemical conditions.** Box plot showing similar surface temperature, salinity, total dissolved solid (TDS), and chlorophyll across all time points. In contrast, a consistent dissolved oxygen (DO) and pH gradient was observed across different months/seasons.

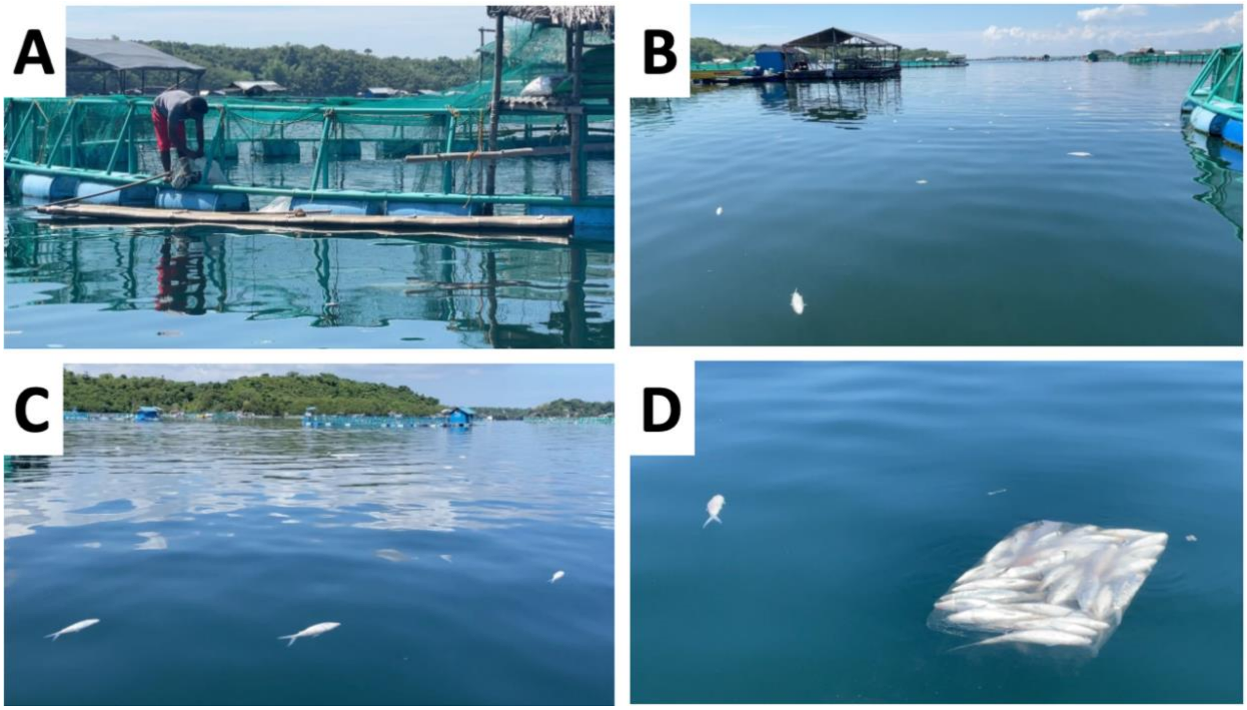


**Figure S3. Fish kill around Station 1 CCMS.** Photos taken May 16, 2022 (A) One of the mariculture caretakers is harvesting dead fish for disposal (B & C). Dead milkfish in the vicinity of Station 1 CCMS. (D) Bagged dead fish for incineration or dumping.

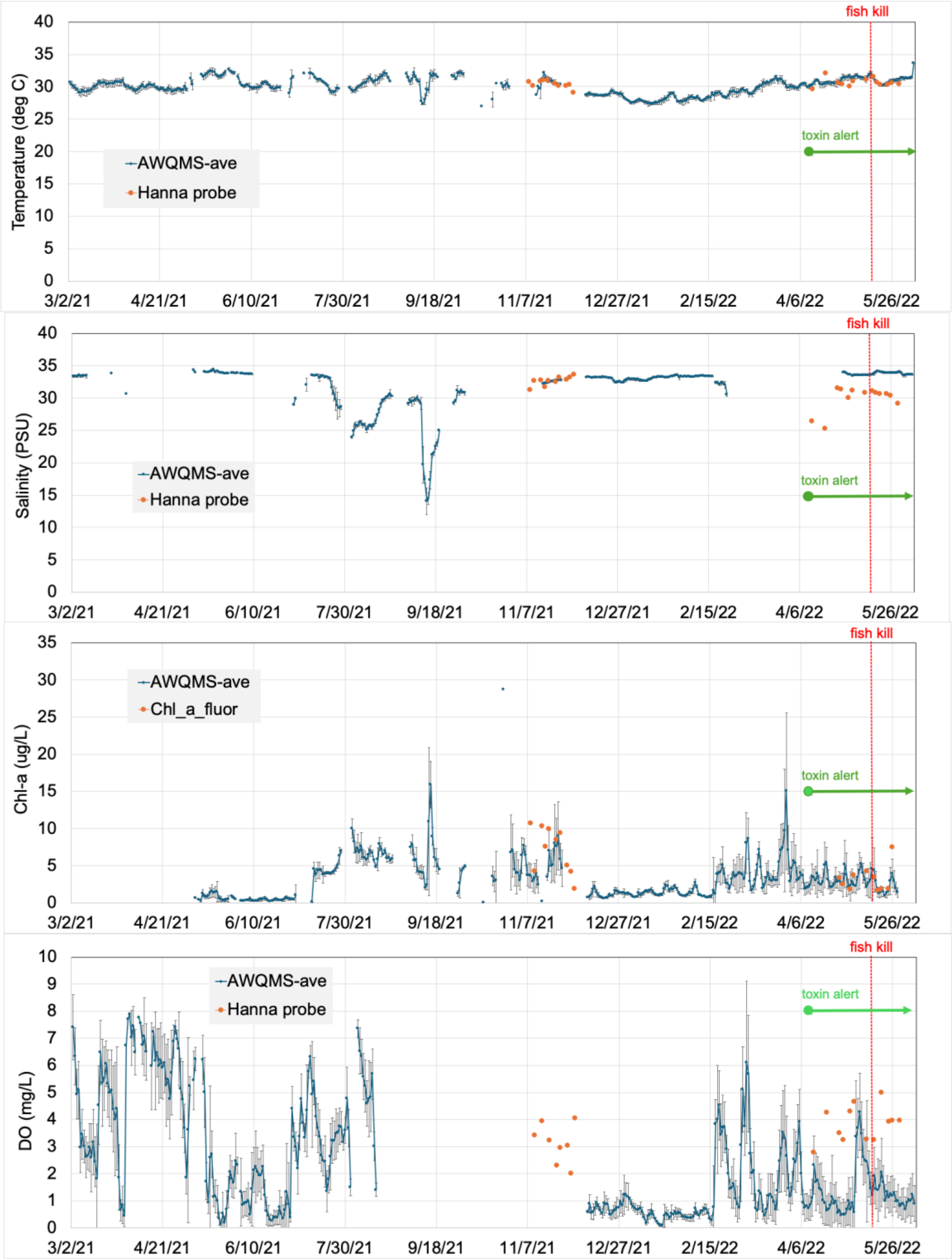


**Figure S4. CCMS data.** In dark blue are the daily averages of AQWMS measurements (with standard deviation). Plotted together are single-point measurements using a Hanna probe (in orange) or lab fluorometer for chlorophyll-a (also in orange). A red line indicates the fish kill event, while a green line indicates the saxitoxin alert.

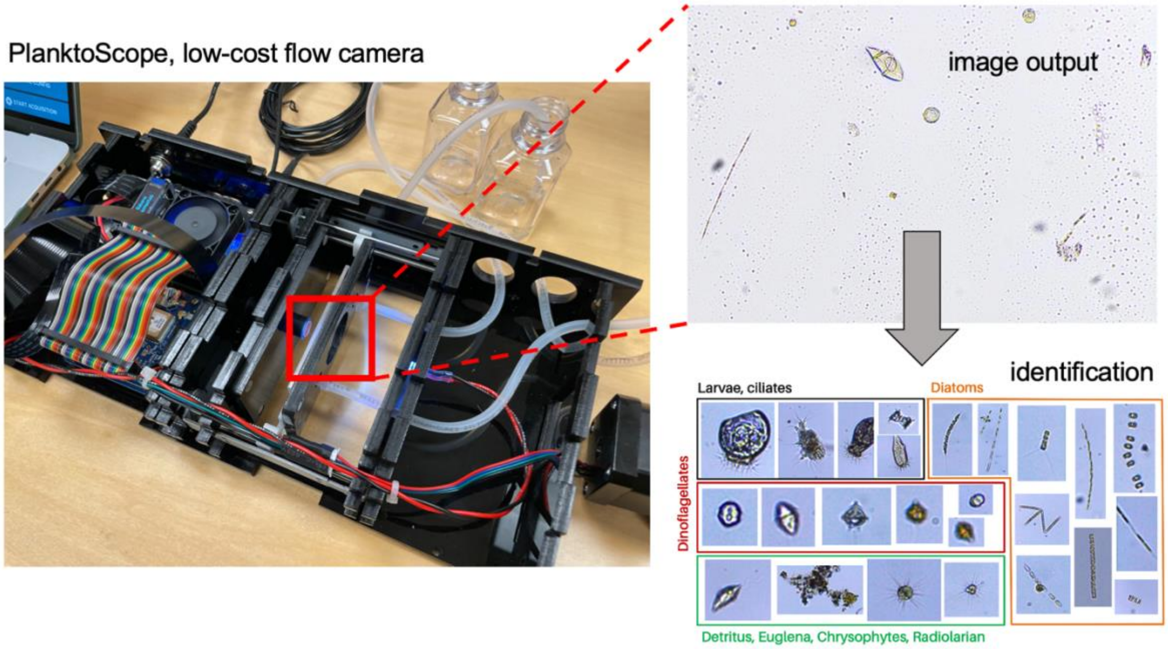


**Figure S5. PlanktoScope v2.1.** Using commonly available parts, a PlanktoScope, developed by the Prakash Lab at Stanford, was replicated and deployed in Bolinao.

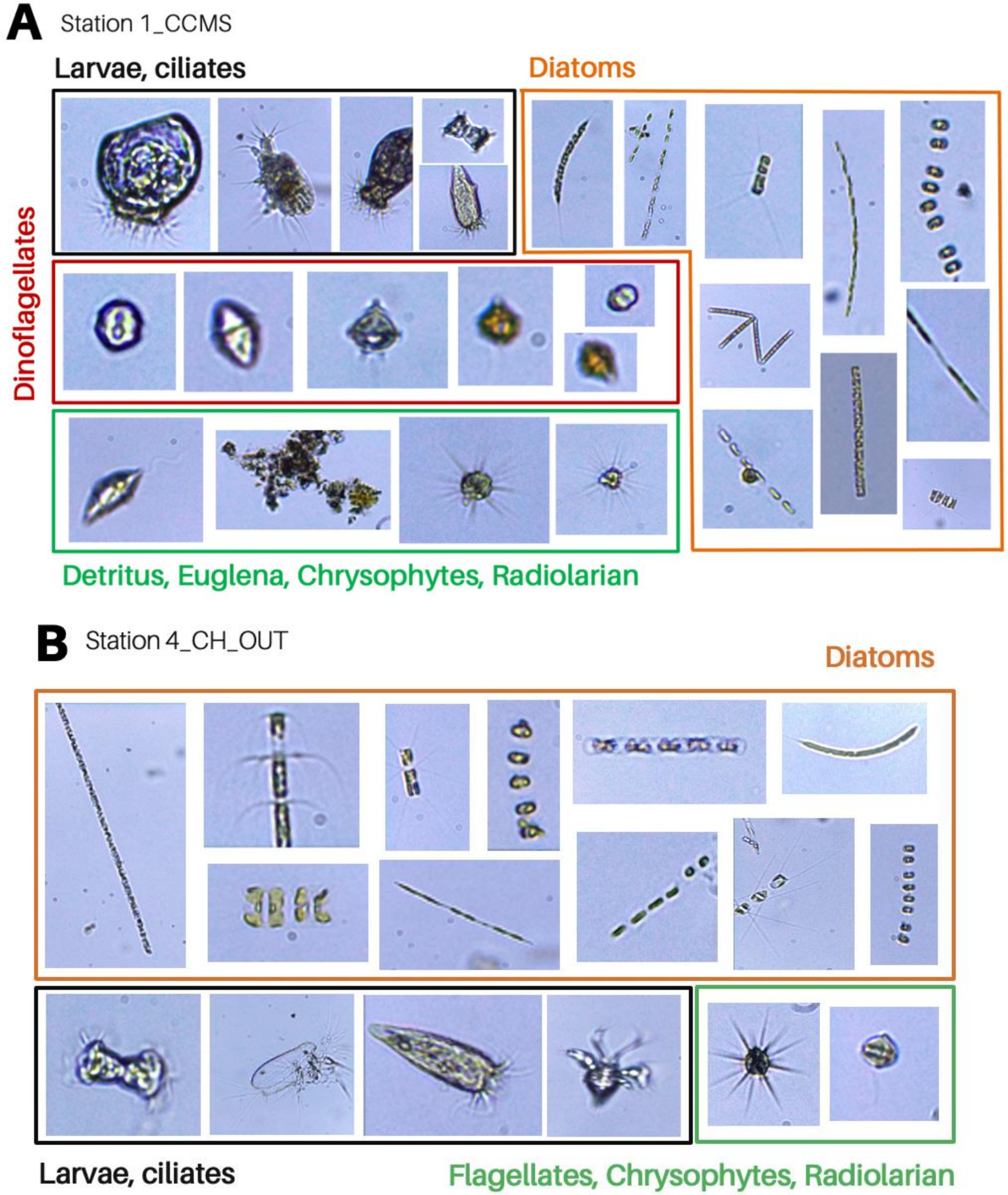


**Figure S6. Representative images from PlanktoScope.** PlanktoScope output for [A] Station 1 CCMS and [B] Station 4 CH_OUT during the Nov-Dec 2021 field sampling.

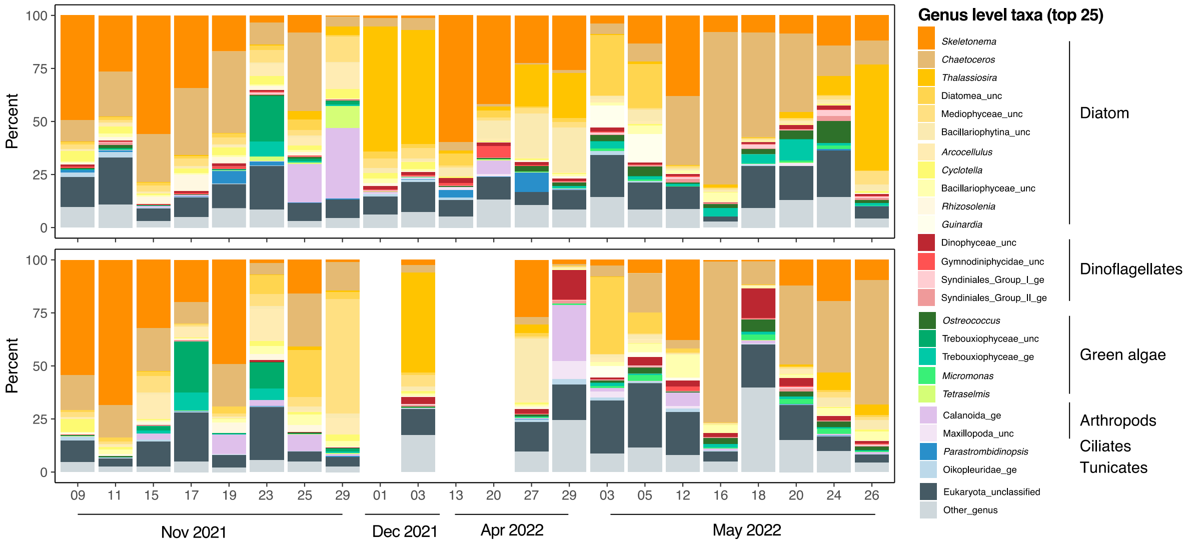


**Figure S7. Plankton relative abundance based on 18S rRNA gene amplicon data (at the genus level, top 25)**

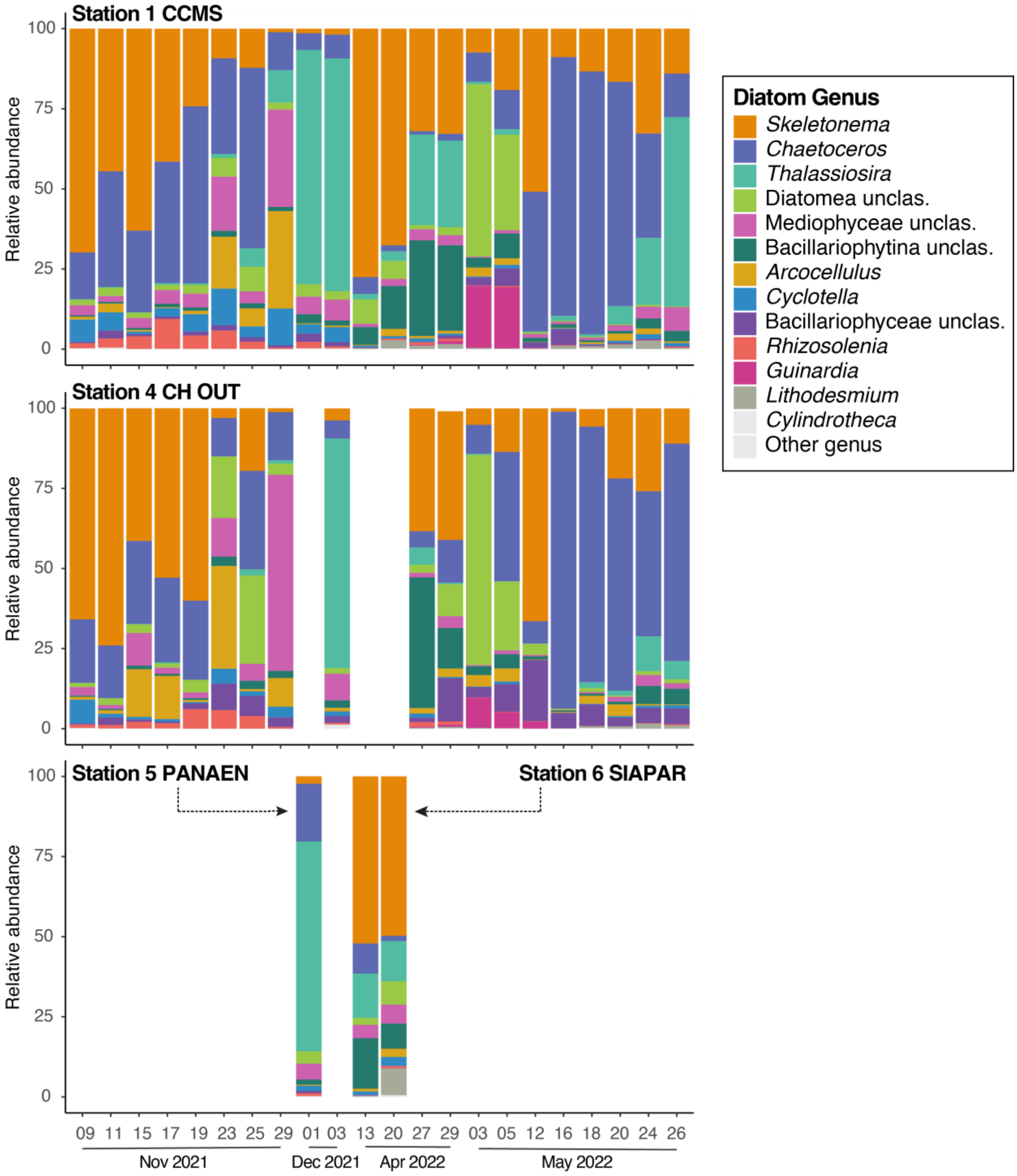


**Figure S8. Relative abundance of diatoms based on 18S rRNA gene amplicon sequences.**

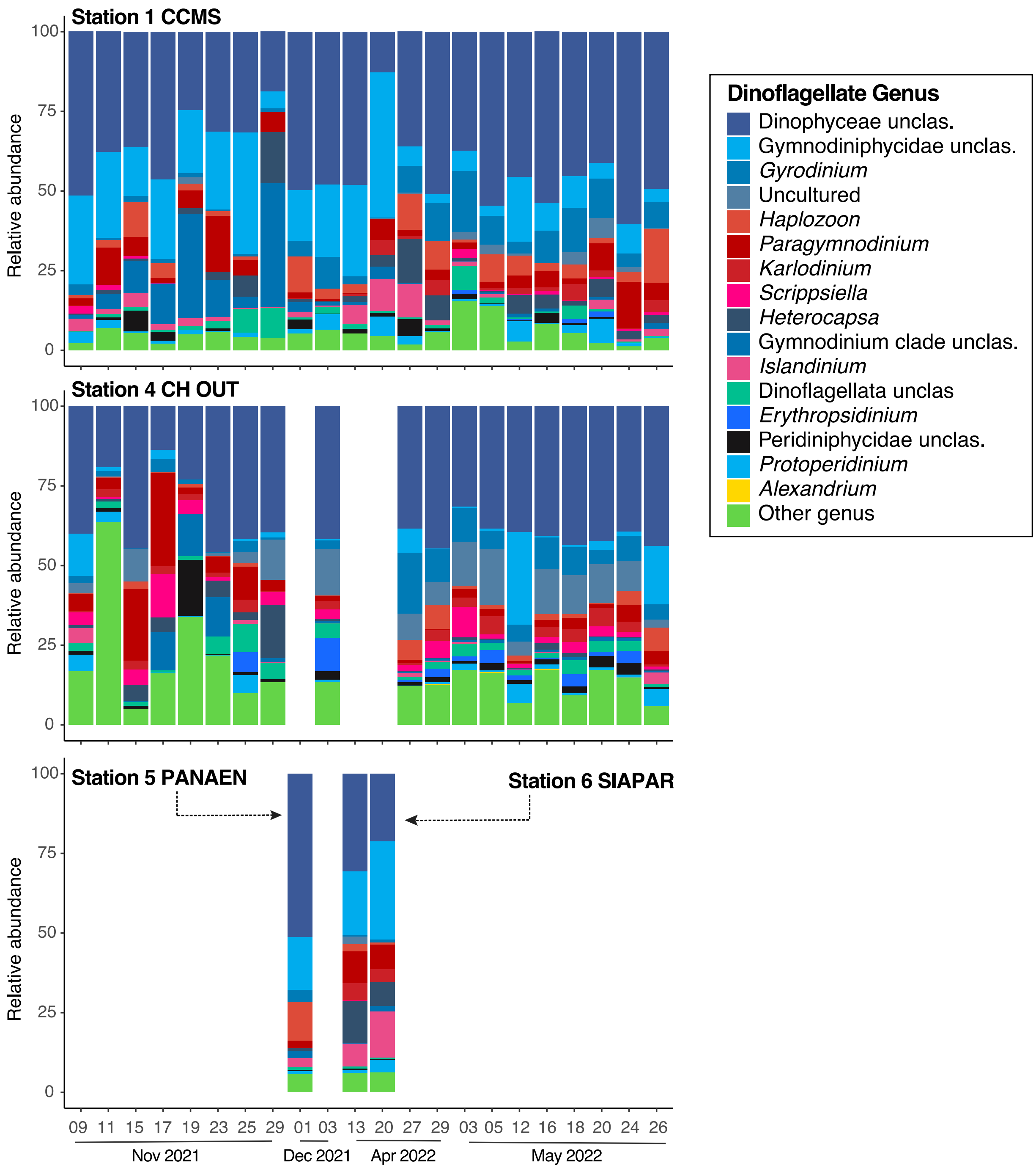


**Figure S9. Relative abundance of dinoflagellates based on 18S rRNA gene amplicon sequences.**

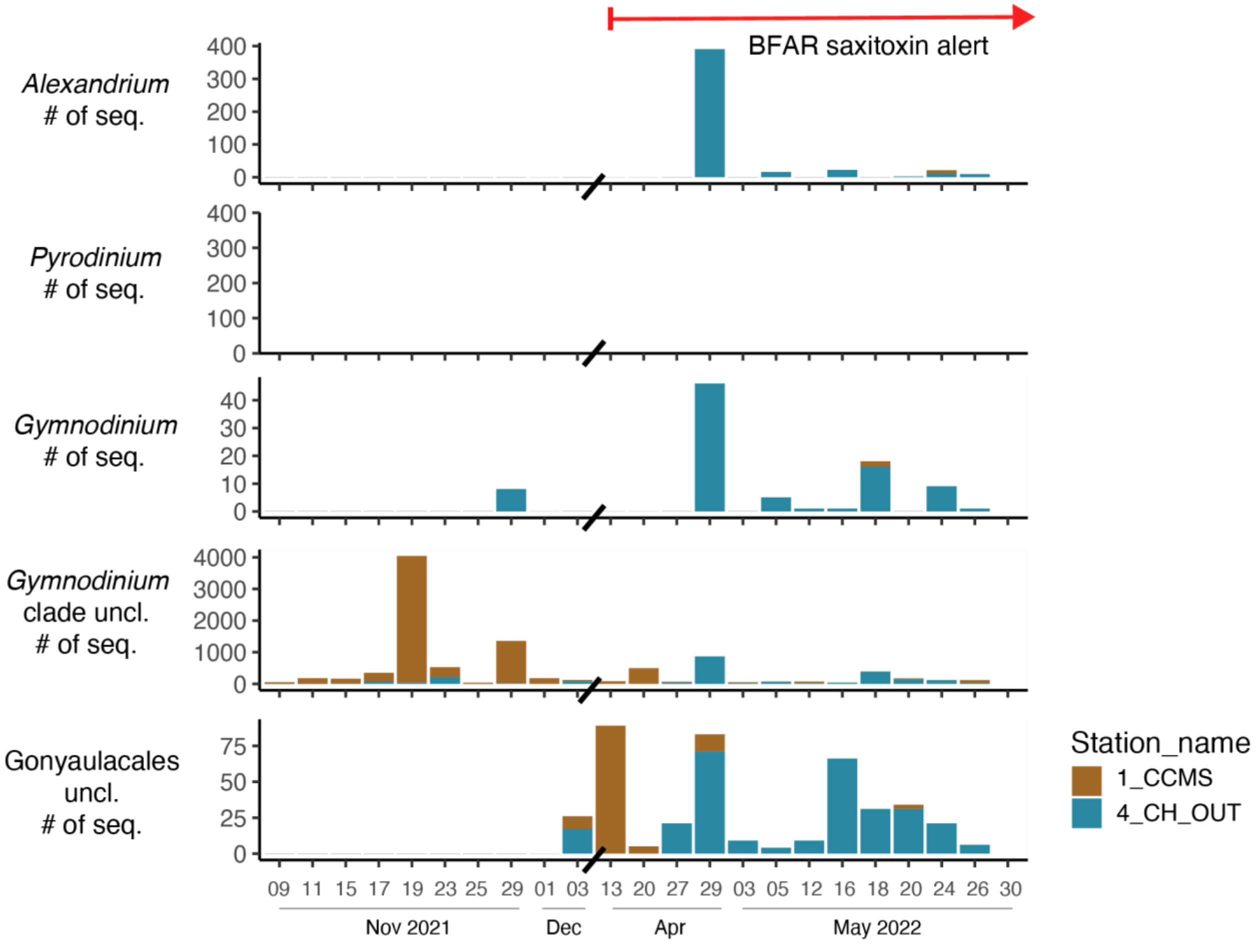


**Figure S10. Relative abundance of potentially saxitoxin-producing dinoflagellates based on 18S rRNA gene amplicon sequences.**

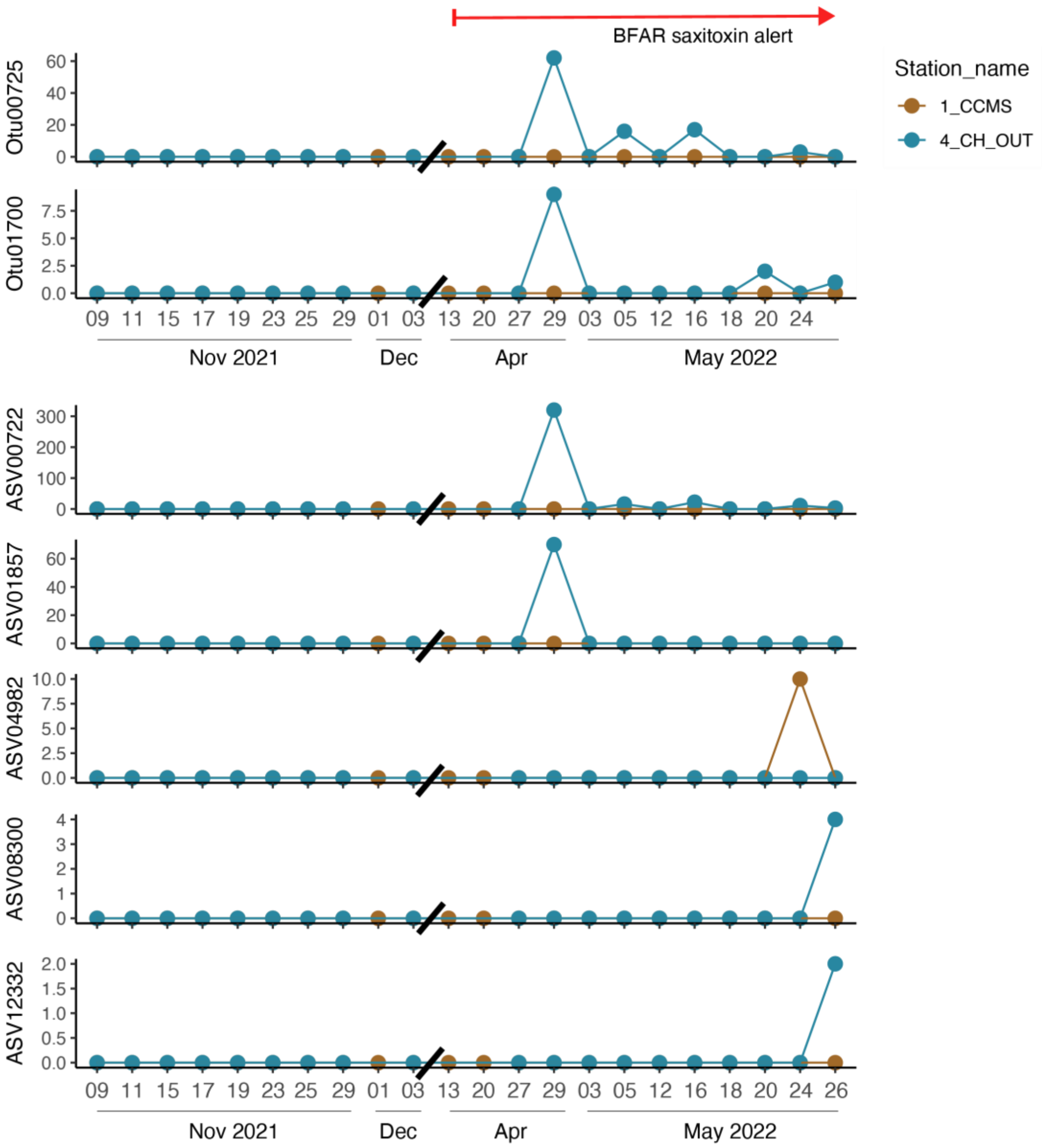


**Figure S11. Temporal dynamics of Alexandrium OTUs and ASVs**

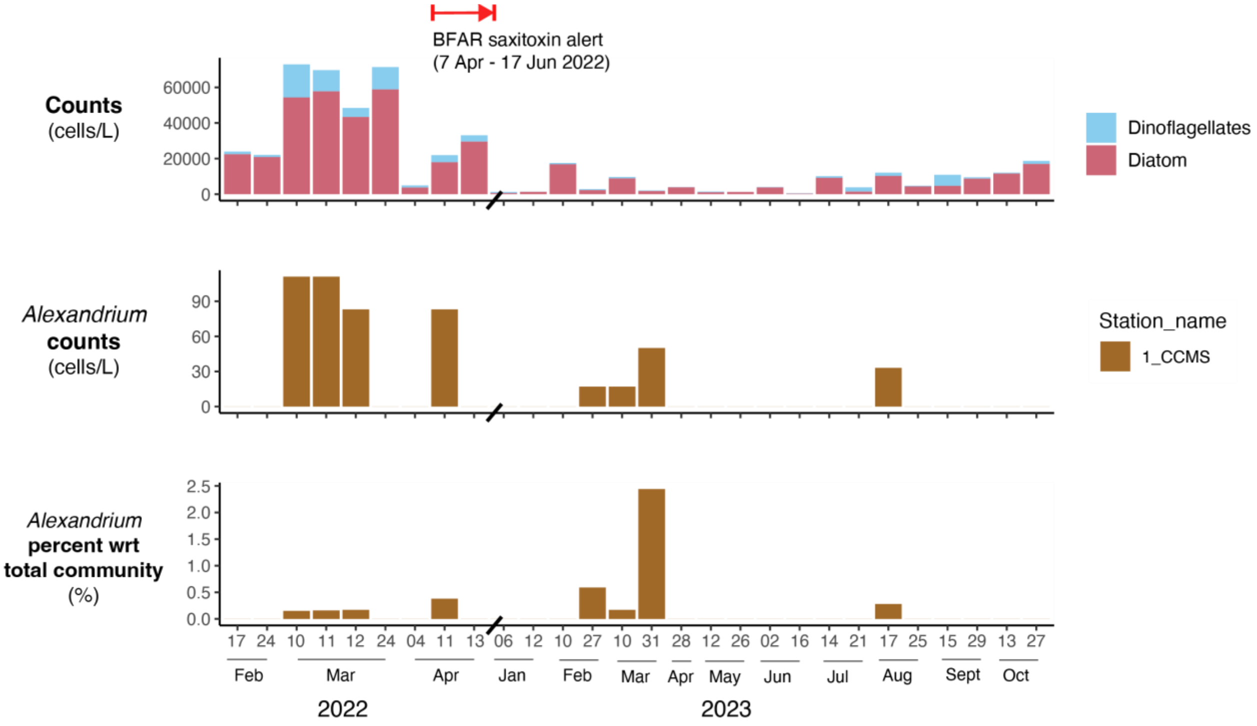


**Figure S12. Phytoplankton abundance and *Alexandrium* counts in CCMS in 2022 and 2023**

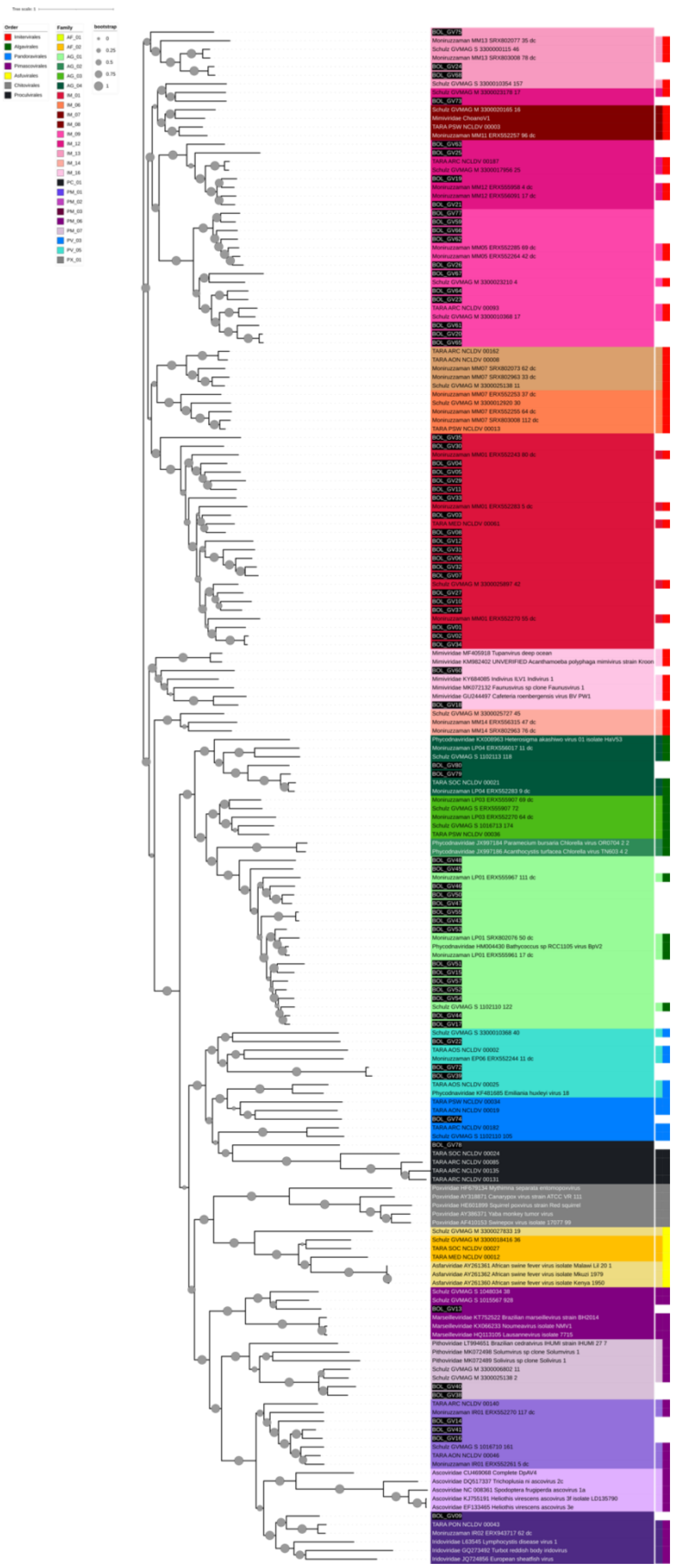

**Figure S13. Phylogeny of Bolinao giant viruses based on the polB gene.**

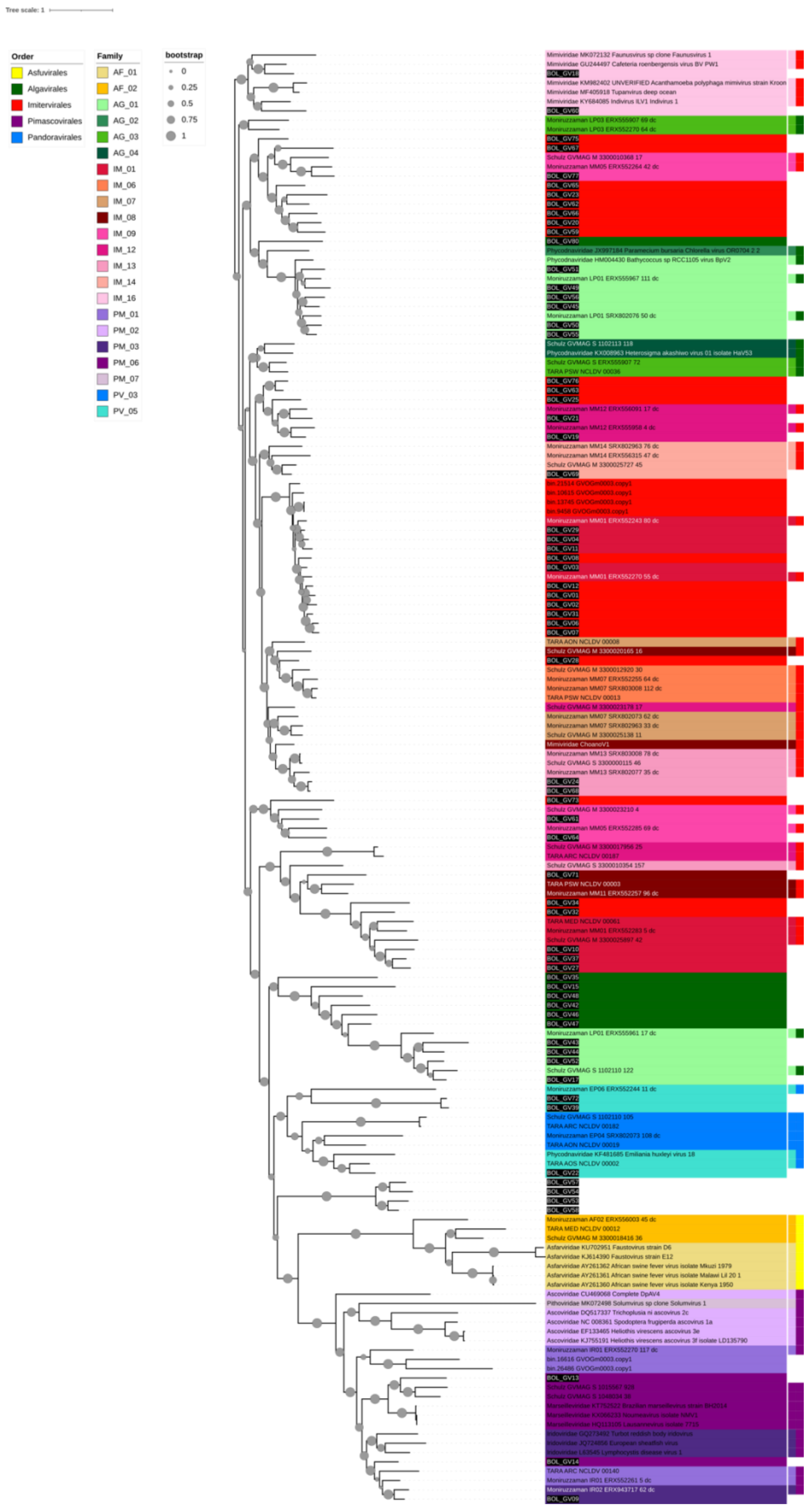

**Figure S14. Phylogeny of Bolinao giant viruses based on MCP gene.**

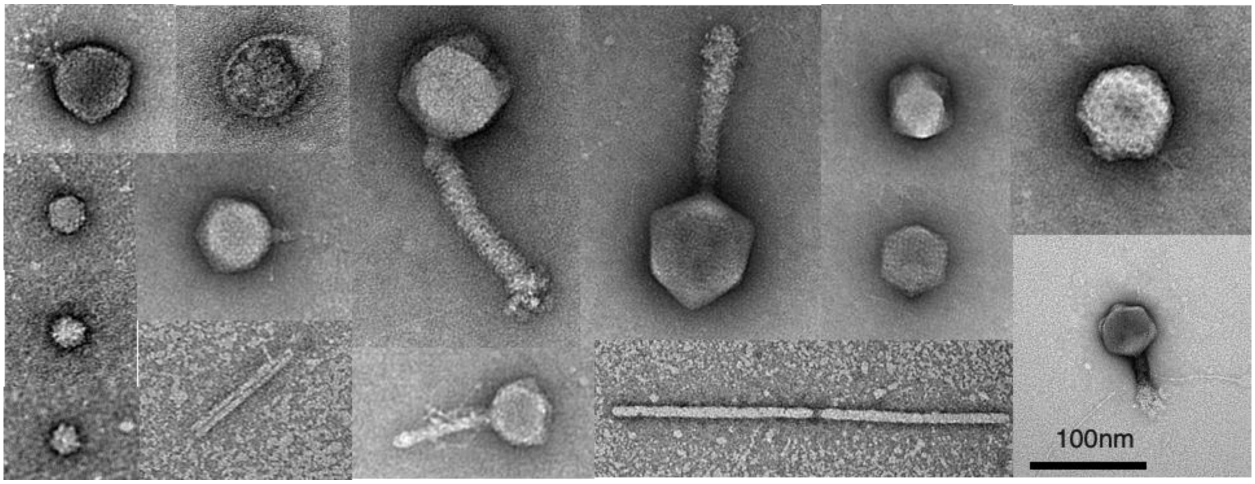


**Figure S15. Other viral-like particles (VLPs) in Bolinao waters.**

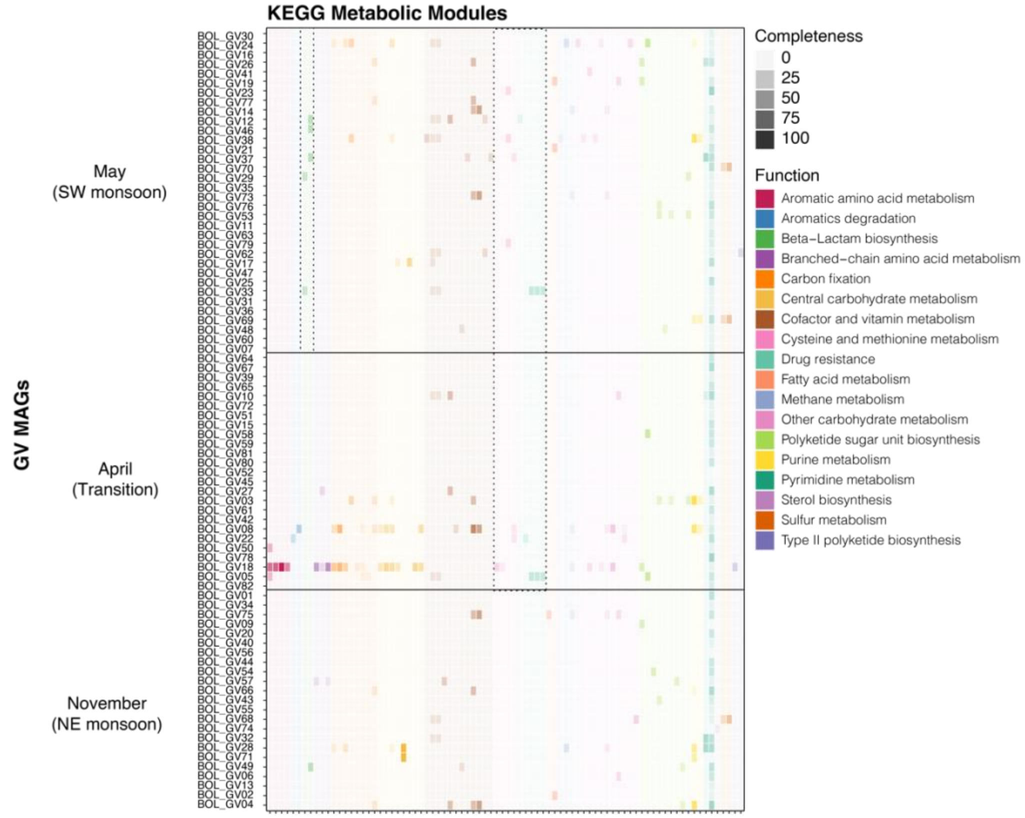


**Figure S16. Bolinao GV MAGs show diverse functional capabilities.** Metabolic capacity was estimated using the KEGG metabolic module completeness. Clustering of GV MAGs is based on season (Figure 6A)

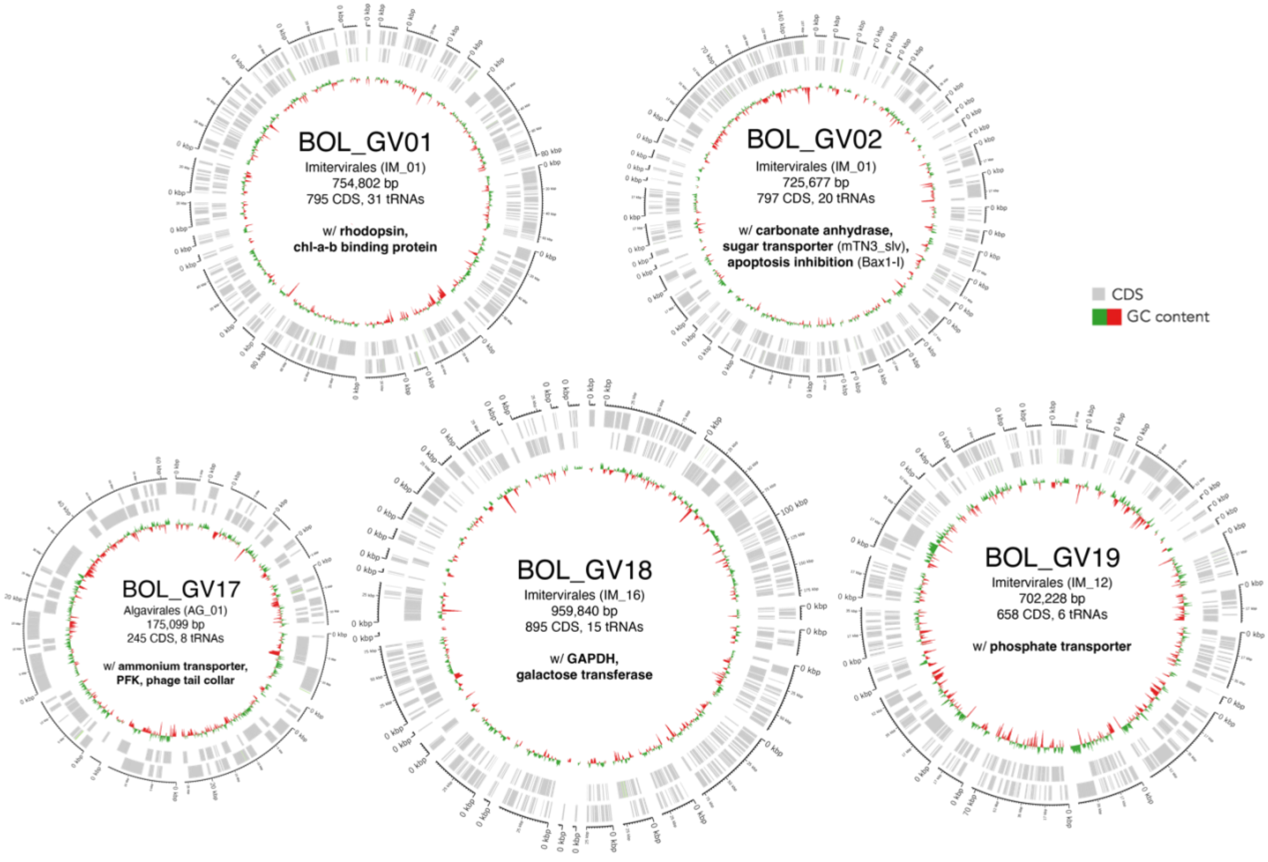


**Figure S17. Genomes of select Bolinao giant viruses.** Bolinao giant viruses encode auxiliary metabolic genes that may confer virocell advantage in eutrophic environments. bp = base pair; CDS = coding sequences

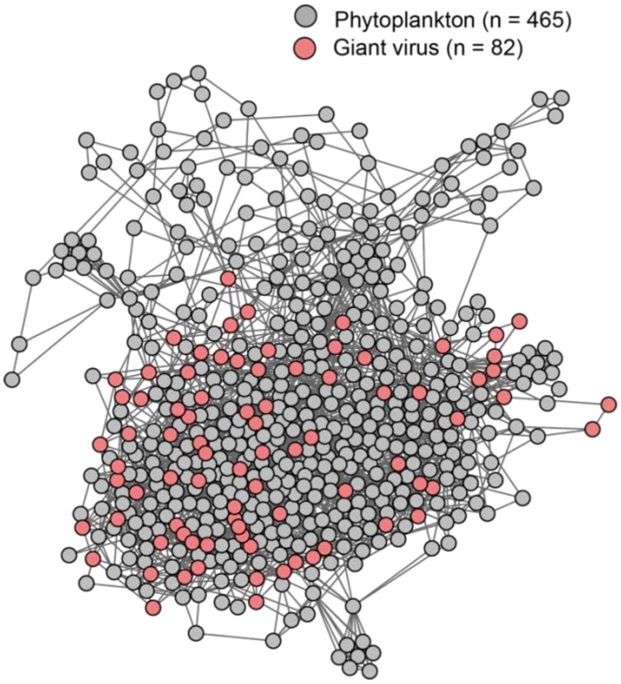


**Figure S18. Co-occurrence network reveals potential phytoplankton-giant virus associations.** After removing diatom and zooplankton OTUs, a co-occurrence network of the remaining plankton and giant viruses was made using SpiecEasi in R. Plotted using iGraph graphopt layout.

**Table S1.** High-quality sequencing reads. Some samples were sequenced twice, corresponding to batches 1 and 2.

| | Sample ID | batch 1 | batch 2 | total |
|---|---|---|---|---|
| 1 | 5-S1-11-11-21 | 40,663,305 | 63,486,455 | 104,149,760 |
| 2 | 8-S4-11-11-21 | 42,253,969 | 60,009,136 | 102,263,105 |
| 3 | 21-S1-11-23-21 | - | 46,056,432 | 46,056,432 |
| 4 | 24-S4-11-23-21 | - | 47,093,621 | 47,093,621 |
| 5 | 40-S1-04-13-22 | 46,753,380 | 136,557,364 | 183,310,744 |
| 6 | 41-S6-04-13-22 | - | 49,134,392 | 49,134,392 |
| 7 | 42-S1-04-20-22 | 49,209,676 | 149,309,607 | 198,519,283 |
| 8 | 43-S6-04-20-22 | - | 50,199,078 | 50,199,078 |
| 9 | 64-S1-05-16-22 | 43,565,697 | 177,791,512 | 221,357,209 |
| 10 | 67-S4-05-16-22 | 59,511,175 | 70,612,972 | 130,124,147 |
| 11 | 76-S1-05-24-22 | - | 54,237,658 | 54,237,658 |
| 12 | 79-S4-05-24-22 | - | 45,252,296 | 45,252,296 |
| 13 | 81-S1-05-26-22 | 52,925,763 | 150,209,980 | 203,135,743 |
| 14 | 84-S4-05-26-22 | 53,767,363 | 53,120,866 | 106,888,229 |
| | | | | **1,541,721,697** |

**Table S2.** Number of reads before and after normalization. Each set was then assembled using MEGAHIT.

| | combined reads | reads after bbnorm |
|---|---|---|
| Station 1_nov | 150,206,192 | 108,142,820 |
| Station 1_apr | 381,830,027 | 255,671,536 |
| Station 1_may | 478,730,610 | 298,980,865 |
| Station 4 | 431,621,398 | 319,559,149 |
| Station 6 | 99,333,470 | 78,264,386 |
| | **1,541,721,697** | **1,060,618,757** |

**Table S3.** Community composition in Station 1 CCMS and Station 4 CH_OUT based on microscopic analysis (counts expressed as cells/L) during the May 2022 sampling.

| | 1_CCMS | | | | 4_CH_OUT | | | |
|---|---|---|---|---|---|---|---|---|
| | 20220516 | 20220524 | 20220526 | 20220530 | 20220516 | 20220524 | 20220526 | 20220530 |
| *Akashiwo* | | | | | | 0.3 | | |
| *Alexandrium* | | | | | | | 0.6 | |
| *Amylax* | 1.4 | | | | 1.3 | | | |
| *Asterionellopsis* | | | | | | | | 0.2 |
| *Bacteriastrum* | | | | | | 4.3 | 10.6 | |
| Bivalve larvae | 2.0 | 3.3 | 1.9 | | 1.9 | | 0.6 | |
| *Ceratium* | | | | | | 0.7 | 0.3 | 0.2 |
| *Chaetoceros* | 32.1 | 6.3 | 6.8 | 2.3 | 29.6 | 45.7 | 52.0 | 22.2 |
| Ciliates | 3.2 | 1.7 | 1.2 | 0.2 | 2.9 | 0.7 | 0.3 | 0.2 |
| Cyanobacteria | | | | | | | 4.4 | |
| *Cymbella* | | | | | | 0.3 | | |
| *Dactyliosolen* | | | | 1.4 | | | 0.6 | |
| *Dinophysis* | | | | | | | 0.3 | |
| *Gonyaulax* | 3.8 | 0.3 | | 0.7 | 3.5 | 1.3 | 1.2 | |
| *Guinardia* | | 0.7 | | | | | | 2.3 |
| *Gymnodinium* | | | 1.2 | | | | 0.3 | 0.2 |
| *Hemiaulus* | | | | 0.9 | | 1.3 | 1.2 | 0.9 |
| *Leptocylindrus* | 1.4 | | 1.2 | 5.5 | 1.3 | 3.7 | 34.8 | 22.4 |
| *Nauplius* | 0.3 | 1.7 | 0.6 | 0.2 | 4.3 | | 0.6 | 0.5 |
| *Navicula* | | 0.3 | 0.6 | 0.2 | | | | 0.5 |
| *Nitzschia* | | 1.0 | 8.4 | 3.5 | | 7.3 | 14.3 | 7.4 |
| *Oikopleura* | | | | | | | 0.3 | |
| *Peridinium* | 6.9 | | 0.3 | | 6.4 | | 0.9 | |
| *Pleurosigma* | 0.3 | | | | 1.3 | | | |
| *Protoperidinium* | 2.0 | 1.0 | 0.3 | 0.2 | 1.9 | | 2.8 | |
| *Pseudo-nitzschia* | 57.8 | 9.7 | 62.2 | 30.5 | 53.3 | 66.7 | 62.2 | 46.2 |
| *Radiolarian* | | | | | | | | 0.5 |
| *Rhizosolenia* | 4.3 | 0.3 | 14.0 | 8.1 | 4.0 | 2.0 | 17.4 | 12.9 |
| *Scrippsiella* | 0.6 | | | | 0.3 | | | |
| *Skeletonema* | 57.8 | 25.7 | 56.9 | 1.4 | 53.3 | 66.7 | | 8.1 |
| *Thalassiosira* | 4.6 | 22.7 | 62.2 | 3.7 | 0.5 | 20.7 | 51.6 | 0.5 |
| Tintinnid | 1.4 | 2.0 | 1.6 | 1.4 | 1.3 | 1.3 | 9.6 | |
